# A standardized bath challenge of Atlantic salmon reveals distinct infection dynamics and mortality across ten HPR-deleted infectious salmon anaemia virus isolates

**DOI:** 10.64898/2026.06.10.731357

**Authors:** Sonal Jayesh Patel, Ole Bendik Dale, Bjørn Spilsberg, Johanna Hol Fosse, Torfinn Moldal, Magnus Leithaug, Marit Måsøy Amundsen, Saima Nasrin Mohammad, Adriana Magalhaes Santos Andresen, Laura V. Solarte-Murillo, Frieda Betty Ploss, Henriette Kvalvik, Simon Chioma Weli

**Author notes:** Corresponding author: Simon Chioma Weli. Aquatic Pathobiology Laboratory, Department of Infectious Diseases and Immunology, University of Florida, Gainesville, FL 32610, USA.

## Abstract

Infectious salmon anaemia virus with highly polymorphic region deletions (ISAV-HPRΔ) is classified as pathogenic, yet field outbreaks display wide variation in disease severity. To determine the extent of inherent virulence differences among ISAV-HPRΔ isolates, we conducted a standardized freshwater bath challenge in Atlantic salmon using ten isolates, including the high-virulent reference strain NO/Glesvaer/2/90 and nine recent Norwegian field isolates. Cumulative mortality, infection kinetics, tissue viral loads, shedding, and pathological changes were characterised through RT-qPCR, histopathology, immunohistochemistry, and flow cytometry. All isolates established systemic infection, but exhibited pronounced differences in infection dynamics, virus shedding, clinical signs, and pathological outcomes. Cumulative mortality ranged from 15% to 100%, allowing separation of isolates into high- (≥90%), moderate- (40–50%), and low-mortality (<20%) categories. Isolates with high mortality showed rapid systemic spread, extensive endothelial infection, and significant pathology compatible with infectious salmon anaemia. Shedding profiles of virus to water differed substantially and were not clearly correlated with cumulative mortality, viral RNA load in tissues or mortality. High ISAV RNA was detected in water for the H16 isolate with ∼10 – 100-fold higher viral RNA than H20 and Å. VA and S, although giving high mortality (>90%), had much lower (shedding (highest RNA range 1.1 – 3.6^10^2^). Segment 5 and 6 sequencing confirmed that all isolates carried genetic mutations typical of pathogenic ISAV except Å, that have an atypical mutation in the putative protease cleavage site on segment 5. However, these mutations alone did not account for the wide biological continuum of mortality.

## Introduction

Viral pathogens remain a leading cause of mortality in aquaculture industry, posing major challenges to fish health management and biosecurity. Among these, infectious salmon anaemia virus (ISAV) is one of the most impactful fish viruses representing a significant threat to the global Atlantic salmon aquaculture industry. ISAV-HPRΔ, the pathogenic variant of the virus, can cause severe disease characterized by anaemia, circulatory disturbances, and multi-organ bleeding, which if allowed to progress can induce substantial cumulative mortality (1, 2). ISA outbreaks have been reported in Norway (3, 4), Chile (5), Canada (6), USA (7), Scotland (8), Iceland (9), and the Faroe Islands (10), leading to vast economic losses and culling of hundreds of millions of Atlantic salmon (11). In Norway, the implementation of strict biosecurity measures have reduced the number of yearly ISA outbreaks from more than 80 widespread outbreaks in 1990 to between one and 25 yearly outbreaks in the last two decades (12).

The current designation of ISAV as pathogenic by the World Organization for Animal Health (WOAH) is primarily based on the presence of a deletion of variable length in the haemagglutinin esterase (HE) highly polymorphic region (HPR) (13). However, this trait does not reflect the full range of biological variations observed among field isolates during outbreaks. Differences between ISAV isolates have been observed in mortality rates, infection dynamics, tissue tropism, and shedding patterns, suggesting that some ISAV-HPRΔ isolates may display lower virulence, reflecting a spectrum within the ISAV-HPRΔ category (14, 15). In addition, a few published experimental studies strongly support that ISAV-HPRΔ isolates may vary considerably in virulence and mortality (14, 16). However, in complex production systems, environmental and host factors can either exacerbate or mitigate the disease outcomes and confound virus-specific contributions to the observed disease severity.

The consequences of not being able to measure these differences in virulence are significant. Management responses against ISA outbreak in Atlantic salmon involve severe measures such as large-scale culling and stringent movement restrictions, which carry economic, social, and ethical costs (17, 18). The current regulation of measures against ISA in Norway are the same for all ISAV-HPRΔ isolates. However, maintaining ISA control by fallowing and surveillance of potential spread is expensive (7). It has been argued that ISA outbreaks in recent decades have been less severe than those in the 1990s. This argument has fuelled assertions that currently circulating ISAV-HPRΔ have low virulence. Based on this reasoning, the industry has expressed a need for “targeted management” approaches to limit culling in cases associated with low mortality. However, using fish mortality in field conditions as a proxy for low virulence and, furthermore, estimation of virulence to determine the implementation of biosecurity measures is rather problematic. Such an approach risks failing to contain virulent viruses whose pathogenic potential is not fully expressed in a given field setting. This is particularly relevant for ISA, where mortality may initially progress slowly but ultimately reach high cumulative levels.

Characterization of ISAV isolates is also useful for vaccination strategies against ISA that has been increasingly employed in several salmon farms in Norway in recent years (19). However, vaccine strategies face several challenges, including difficulties in selecting vaccine strain that may offer protection against most ISAV isolates, challenges in monitoring ISAV evolution that may diminish the effectiveness of existing vaccines, differentiating vaccinated from infected salmon, and maintaining appropriate vaccination coverage in the ISA hotspot zones.

In this study, we investigate potential inherent differences in virulence by conducting a standardized experimental infection in Atlantic salmon with a panel of ISAV field isolates. The ISAV panel included a type-strain (Norwegian isolate NO/Glesvaer/2/90) with well-characterized high virulence, and nine ISAV-HPRΔ field isolates collected from more recent outbreaks, including three newly emergent isolates with no or very limited clinical signs of ISA disease, and one isolate with an atypical point mutation in the F protein cleavage site region. To generate detailed comparative profiles for each isolate, we analysed infection dynamics, viral shedding, pathology, mortality, and genome sequences.

## Materials and Methods

### Sequencing and choice of isolates

Samples from the last decade, hereafter called recent ISA outbreaks were selected for RNA isolation, amplicon library preparation, and whole genome sequencing was performed as described by Spilsberg et al. (20). Based on a phylogenetic tree containing 83 isolates (result not shown), isolates for the challenge trial were selected representing different genetic clusters also ensuring geographical spread (Table 1). The well-characterised type-strain isolate NO/Glesvaer/2/90 was included as positive control. Read counts and quality were recorded with FastQC version 0.12.1 (21) and MultiQC version 1.29 (22). Gene specific primers were removed with BBduk from the BBmap package version 39.37 (23) with the following parameters: remove last base, trim in the first 30 bases, with k=17, edit distance=2 and quality trimming at phred score=15. Illumina adapters were removed with Trimmomatic version 0.39 (24). Finally, the forward and reverse read were merged with NGmerge version 0.4 (25). Each quality trimmed read was mapped with Bowtie2 (26) to HQ259675.1 for segment 5 and HQ259676.1 for segment 6, respectively. Consensus sequences were called from the mapped read files with iVar version 1.4.4 (27) using a minimum depth of 10. Each segment was aligned with Muscle (28) in MEGA12 (29) and the evolutionary distances were calculated using a Kimute 2 parameter model (30). Multiple sequence alignments (MSA) were visualized in R version 4.4.1 using the packages Tidyverse and GGMSA.

**Table 1.** Isolates used in this study and a HPR0 reference strain (SK) used for sequence comparison.

| Isolate acronym | Isolate name | Aquaculture facility (number) | Isolation year | Accession segment 5 (F) | Accession segment 6 (HE) |
| --- | --- | --- | --- | --- | --- |
| GL | Glesvaer/2/90 | Unknown | 1990 | HQ259675.1 | HQ259676.1 |
| VE | NO/Heim/NVI-70-122/2022 | Vestseøra (24096) | 2022 | PZ269547.1 | PZ269548.1 |
| VA | NO/Lofoten/NVI-70-2774/2014 | Våtvika (13047) | 2014 | OQ310960.1 | OQ310972.1 |
| HG | NO/Heim/NVI-70-150/2022 | Hønsvikgulen (12898) | 2022 | PZ269549.1 | PZ269550.1 |
| H16 | NO/Salten/NVI-70-1297/2016 | Hestholmen (13006) | 2016 | PZ269551.1 | PZ269552.1 |
| H20 | NO/Salten/NVI-70-595/2020 | Hestholmen (13006) | 2020 | PZ269553.1 | PZ269554.1 |
| S | NO/Sor-Troms/NVI-70-663/2021 | Salangsverket (36357) | 2021 | PZ269555.1 | PZ269556.1 |
| Å | NO/Helgeland/NVI-70-1169/2015 | Åmøya (26375) | 2015 | PZ269557.1 | PZ269558.1 |
| B | NO/Vesteralen/NVI-70-9227/2013 | Bonhammaren (11248) | 2013 | OQ310956.1 | OQ310968.1 |
| L | NO/Finnmark/NVI-70-72/2020 | Laholmen (15460) | 2020 | PZ269545.1 | PZ269546.1 |
| SK* | SK779/06 | Unknown | 2006 | EU118819.1 | EU118820.1 |

\*Reference HPR0 virus, not included in bath challenge trial

### Cell culture

Atlantic salmon kidney (ASK) cells (ATCC® CRL-2747™) (31) were maintained at 20°C in closed cap T-175 tissue culture flasks containing L-15 medium (Biowest) supplemented with foetal bovine serum (FBS,10%, Avantor), L-glutamine (VWR, 4mM), and penicillin/streptomycin/amphotericin (Biowest, 1%).

### Virus propagation and titre determination

The highly virulent Norwegian ISAV isolate NO/Glesvaer/2/90 (32) (GL) and nine other recent ISAV isolates (VE, VA, HG, H16, H20, S, Å, B and L, Table 1), were propagated in ASK cells as previously described (32). Briefly, ASK cells were infected with passage 2 viruses and incubated at 15 °C, and supernatants were harvested when cytopathic effects were close to complete, 9-15 days post infection (DPI). Harvested supernatants were clarified by centrifugation (3800 ×*g*, 10 min, 4 °C). Infective titres were determined by inoculating serial dilutions of supernatants in six parallel wells of ASK cells cultured in 96-well microtiter plates. The plates were incubated at 15 °C for 7 days. Acetone-fixed cells were incubated with IgG1 against the ISAV nucleoprotein (P10, Aquatic Diagnostics Ltd, 0.4 μg/mL, 60 min, room temperature, RT), washed (PBS with 0.1% Tween 20 × 3), incubated with biotin-labelled goat anti-mouse Ig antibody (127-10395, Ray Biotech diluted 1:200 in PBS, 60 min, RT), washed 3x, followed by incubation with FITC-labelled streptavidin (11-4317-87 Invitrogen diluted 1:100 in PBS, 60 min, RT). After a final washing (3x), the cell nuclei were stained by adding 50 μL of a 40 μg/mL propidium-iodide solution per well. The plates were incubated for 2 min at RT and washed once before examination under an inverted fluorescence microscope (Nikon Eclipse Ti2). Virus titre was calculated according to the 50% end-point method of Reed and Muench (1938) (33) and expressed as the dilution of the virus causing 50% tissue infection per millilitre (TCID_50_/mL).

### Fish

The experiment was carried out at the experimental animal facility at the Centre for Sustainable Aquaculture at the Norwegian University of Life Sciences (NMBU), Ås, Norway in accordance with current animal welfare regulations. The protocol was pre-approved by the Norwegian Food Safety Authority, FOTS ID 30432. Atlantic salmon fish line of AquaGen Robust Gen-innova Shield line was supplied by the rearing facility at the Centre for Sustainable Aquaculture, NMBU. The fish were screened by the supplier and tested negative for ISAV, salmon gill pox virus (SGPV), infectious pancreatic necrosis virus (IPNV), piscine orthoreovirus-1 (PRV), piscine myocarditis virus (PMCV), and salmonid alphavirus (SAV). Experimental fish were unvaccinated Atlantic salmon post-smolts with an average weight of 50-80 g. The fish were maintained in tanks with flow-through aquaculture system supplied with filtered (20 µm), UV-treated freshwater. A constant temperature of 12 °C, a daily photoperiod of 12:12 h light:dark, and oxygen saturation of about 80% were maintained throughout the experimental trial period.

### Experimental ISAV Bath Challenge

An overview of the bath experimental design is shown in Fig 1. After acclimatization, 110 fish per isolate were infected with ISAV (10^3.5 TCID_50_ per L tank water) in a 2-hour bath challenge as previously described (15) in freshwater at 12 °C. During the challenge, the water flow was stopped, tanks were continually aerated, and the oxygen levels were closely monitored. Ten fish groups were infected with the respective ISAV isolates, including the high virulent historical Norwegian ISAV isolate NO/Glesvaer/2/90 (GL) (32) and nine recent ISAV field isolates (VE, VA, HG, H16, H20, S, Å, B and L, Table 1). After 2 hours, fish infected with each isolate were transferred using a net to avoid transferring virus containing water from the infection tank to maintenance tanks. Two identical 450 L maintenance tanks provided separate fish groups for sampling (SG, n = 70) and observation (OG, n = 40), respectively. After transfer, the fish were monitored for signs of distress for one hour. Infections with isolate B and L were performed 14 days after the other isolates, with 24 and 21 fish in the SG, respectively. For B, there were 40 fish in OG, while for L no OG could be included. The restrain in fish numbers for isolate B and L was due to much lower virus titre attained during propagation. An additional tank with 102 non-infected fish served as controls. The water flow rate in all tanks was adjusted according to biomass to maintain dissolved oxygen level. The challenge experiment lasted for 42 days for all isolates except for isolates B and L for which the duration was 28 days, with daily recording of fish mortality. Several OGs and SGs were terminated earlier upon reaching 100% mortality or when moribund fish combined with sampled fish resulted in SGs had no remaining fish.

**Figure 1.**
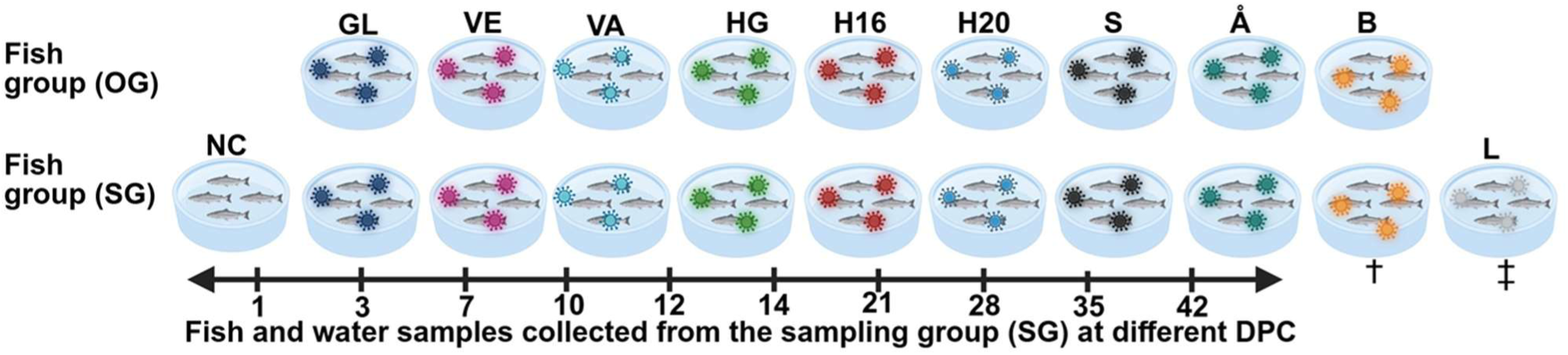
Schematic illustration of bath experimental design with Atlantic salmon and Norwegian isolate NO/Glesvaer/2/90 and nine recent ISAV field isolates (VE, VA, HG, H16, H20, S, Å, B and L, full names shown in Table 1). Note: OG = Observation fish group with n=40 for all isolates except VA and B with *n= 45 and 42 respectively*. SG = Sampling fish group with n=70. †B = In SG, only twenty-four in infection trial, and fish samples were collected on 3, 7, 14 and 21 days post challenge (DPC). ‡L = Only SG group with twenty-one fish in infection trial. OG was not included for L, and fish samples were collected on 3, 14, 21 and 28 DPC. N= 102 in non-infected (NC) SG.

### Sampling

#### Water sampling

Monitoring the amount of viral RNA in wastewater has been shown to be effective for the detection of marine fish viruses (34, 35). The approach, relying on the shedding of virus from infected fish and its subsequent concentration from water before detection of viral RNA by RT-qPCR, was used to assess ISAV shedding into the water in all experimental groups. Water sampling from the SGs for ISAV RNA detection was performed before onset of challenge and at 1, 3, 7, 10, 12, 14, 21, 28, 35, and 42 days post challenge (DPC). For the SGs with high mortality reaching, the groups were terminated when last fish was sampled, and thus water sampling was precluded thereafter. For the L and B groups, sampling period was shorter with fewer timepoints as the trial lasted for maximum 28 days. At each time-point, 2 × 250 mL of water was sampled from each of the SGs. The concentration of the 250 mL water samples was performed as described previously (36), and viral RNA was extracted and stored at −80 °C prior to quantitative (RT-qPCR) analysis.

#### Fish sampling

Four fish from eight of ten isolates in SGs and the non-infected control group were sampled at 1, 3, 7, 10, 12, 14, 17, 21, 28, 35, and 42 DPC, in addition to four fish sampled before challenge (0-samples). For isolate B, fish samples were collected at on days 3, 7, 14 and 21, while for L at 3, 14, 21 and 28 DPC. For the SGs with high mortality, the groups were terminated when last fish was sampled, and thus last sampling time-point and number of fish for GL, VA, and B was 21 DPC (N=2), for S 28 DPC (N=3), for L 28 DPC (N=2), and for Å 35 DPC (N=2). Fish were anesthetized by immersion in benzocaine (40 mg/L), before blood was collected from the caudal vein into heparinized syringes. After blood sampling, necropsy was performed. Gill, heart, and a sub-sample of whole blood were aseptically collected in individual RNA-Later tubes (Qiagen, Hilden, Germany) for RT-qPCR. Gill, pseudobranch, heart, spleen, kidney, liver, and skin/muscle samples were fixed in 10% phosphate buffered formalin for at least 24 hours for histopathology and immunohistochemistry (IHC) before further processing.

### RNA extraction and RT-qPCR

Gill and heart tissue samples (15-30 mm^3^) were collected separately in RNA-later™ (Thermo Fisher Scientific), transferred to 500 μL MagNA Pure LC RNA Isolation Tissue Lysis buffer, and homogenized with 3-5 mm steel beads in a TissueLyser II (Qiagen, 24 Hz, 2 × 3 min). For RNA extraction from blood, 30 μL heparinized whole blood was added to 400 μL MagNA Pure LC RNA Isolation Tissue Lysis buffer (Roche). A total of 350 μL of lysed blood or homogenised tissue (gill or heart) was transferred to each of a Magna Pure 96 Processing Cartridge (Roche) per sample. RNA was extracted on a MagNA Pure 96 instrument (Roche) with the MagNA Pure 96 Cellular RNA Large Volume Kit (Roche), using the RNA tissue FF standard cellular RNA protocol with an elution volume of 50 μL per sample. RNA yield and purity was determined by a Multiskan Sky spectrophotometer (Thermo Fisher Scientific). The primer sequence targeting ISAV segment 8 (forward primer CTACACAGCAGGATGCAGATGT, reverse primer CAGGATGCCGGAAGTCGAT and probe CATCGTCGCTGCAGTTC 5’-6-FAM, MGB-NFQ-3’) was used for amplification of the genetic region encoding the ISAV matrix protein (37). All reactions were performed in 20 μL, containing 5µl of eluted RNA per reaction, 0.4 μM of each primer, 0.2 μM of probe, Brilliant III Ultra-Fast QRT-PCR Master Mix (600884, Agilent Technologies), and nuclease-free water. Cycling conditions on Biorad CFX PCR system were 3 min denaturation at 95 °C, followed by 45 cycles at 95 °C for 5 s and 60 °C for 10 s. Standard curves of a synthetic DNA fragment containing the relevant target sequences (Integrated DNA Technologies) were used for calculation of copy number per PCR reaction. The results were analysed, and graphs were drawn with the help of GraphPad Prism 10 (GraphPad Software Inc., San Diego, CA, USA).

To compare the results between fish in SGs infected with GL and each test isolate, two-way ANOVA with Dunnett’s multiple comparisons test was carried out and presented as heatmaps. Results from RT-qPCR analysis from gills, heart, and whole blood (log_10_-transformed to stabilize variance and allow comparison between groups), in addition to the flow cytometry data were used for this comparison. The analysis excluded (i) isolates B and L due to the low number of sampling time points, (ii) sampling time points with less than four fish per isolate (21 DPC onwards), (iii) sampling time points with no detection of virus in the type strain GL (1DPC heart, 1DPC blood, including flow cytometry).

### Quantification of ISAV- blood cell interactions by flow cytometry

Heparinized blood was stored at 4 °C until immunostaining and analysis. The blood pellet was washed once by the addition of 1 mL PBS and centrifugation (500 ×*g*, 15 °C, 5 min), 10 µL pelleted cells were suspended in 1000 µL PBS, and 100 µL of this suspension was transferred to duplicate wells in a U-bottom 96-well microplate. To remove heparin traces and wash the cells, the plate was centrifuged (500 ×*g*, 15 °C, 5 min), supernatants discarded, and the pellets were resuspended in 50 µL primary antibody (mouse IgG1 anti-ISAV HE, clone 3H6F8, 1/10) (38) or PBS. After incubation (room temperature, 60 min), 100 µL PBS was added to each well before centrifugation and two additional washes, each in 150 µL PBS. The pellet was resuspended with goat anti-mouse IgG (H+L) Alexa Fluor 488 (Molecular Probes, #A11001, 5 ug/mL), incubated at room temperature, shielded from light, for 45 min. Further, cells were washed in PBS as above, resuspended in 100 µL PBS, and analysed by flow cytometry. Gating based on forward scatter (FSC) and side scatter parameters were used to identify cells and exclude debris, and doublets were excluded using FSC-A versus FSC-H gating. Fluorescence thresholds for Alexa Fluor 488 positivity were defined based on unstained and PBS-treated controls to account for autofluorescence and background signal. Fluorescence was measured for at least 20,000 cells (10,000 cells for negative control, PBS-stained wells) using a Novocyte Flow Cytometer (Agilent). To account for potential strain-specific differences in antibody affinity, the signal was expressed as the percentage of ISAV HE-positive cells exceeding the fluorescence range of cells from non-infected fish. Flow cytometry data were analysed using NoVo Express (Agilent) software, and consistent gating strategies were applied across all samples. While our analysis did not distinguish between different blood cell types, we have previously demonstrated that the dynamics of ISAV-blood cell interactions detected by flow cytometric quantification closely mirror the dynamics observed by manual counting of ISAV-positive erythrocytes (RBC) on blood smears from the same fish (39).

### Histopathology and immunohistochemistry

Fifty non-infected control fish and 365 infected fish were processed for histology. Formalin fixed tissue samples were dehydrated and embedded in paraffin. Thin tissue sections (3 μm) on Superfrost slides were prepared for HE-staining, placed on Polysine Adhesion Microscope slides for IHC, heat treated (60 ± 3 °C, minimum 20 min) and deparaffinized before staining. For histological evaluation, first two fish for all isolates were stained with hematoxylin & eosin (H&E), scanned with a Hamamatsu NanoZoomer S360 and read digitally using NDP.2 software. Histopathology scoring of liver and kidney lesions indicative of ISA was performed on 205 infected and 24 control fish sampled from day 7 to day 28. The semiquantitative scores were as follows; not found (0), very sparse (0.5) sparse (1), sparse-moderate (1.5), moderate (2), moderate to severe (2.5), and severe (3). Cellular pathology in the spleen as shown in Fig 11A was scored in the same manner, and average scores for each organ and isolate were calculated (Table 4). The tissue distribution of the viral protein was investigated by IHC for selected isolates resulting in high mortality (GL and VA), moderate mortality (HG), and low mortality (VE). All sampled organs from two fish per isolate, sampled on day 7, 10, 14, and 21 were subjected to IHC to detect ISAV nucleoprotein. Briefly, mouse IgG1 targeting the ISAV nucleoprotein (P10, Aquatic Diagnostics Ltd., Stirling, Scotland) was used for detection of ISAV, employing the Vectastain ABC-AP kit (Vectastain anti-mouse Ig ABC-AP kit; Vector Laboratories Inc.) and Fast Red as the substrate.

## Results

Nine isolates from outbreaks along the Norwegian coast during the last decade were selected for the challenge trial. The selection aimed at capturing both sequence variation and geographical spread. As a positive control, the well characterized pathogenic NO/Glesvaer/2/90 isolate was used. The isolates from V and HG were selected as these most likely originated from the same ISAV HPR0 and only differed in HPR. Two isolates from outbreaks four years apart at the same site (Hestholmen N) were included.

### Onset and Progression of Mortality

All ten ISAV isolates caused lethal infection. Among the nine isolates with OGs, the cumulative mortality ranged from 15% to 100%. In the VA- and GL-infected OGs, the first mortality was observed at 11 and 12 DPC respectively, with cumulative mortalities increasing rapidly, reaching 98%. Isolate B had the highest mortality, reaching 100% by 28 days, while VE gave lowest mortality of 15% (Fig. 2). For the tenth isolate, L, no OG was included in this trial. Mortality was observed in the L-infected SG; however, concurrent sampling in this tank precluded accurate estimation of mortality and direct comparison with the other isolates. Consequently, mortality for L is not shown in Fig. 2.

**Figure 2.**
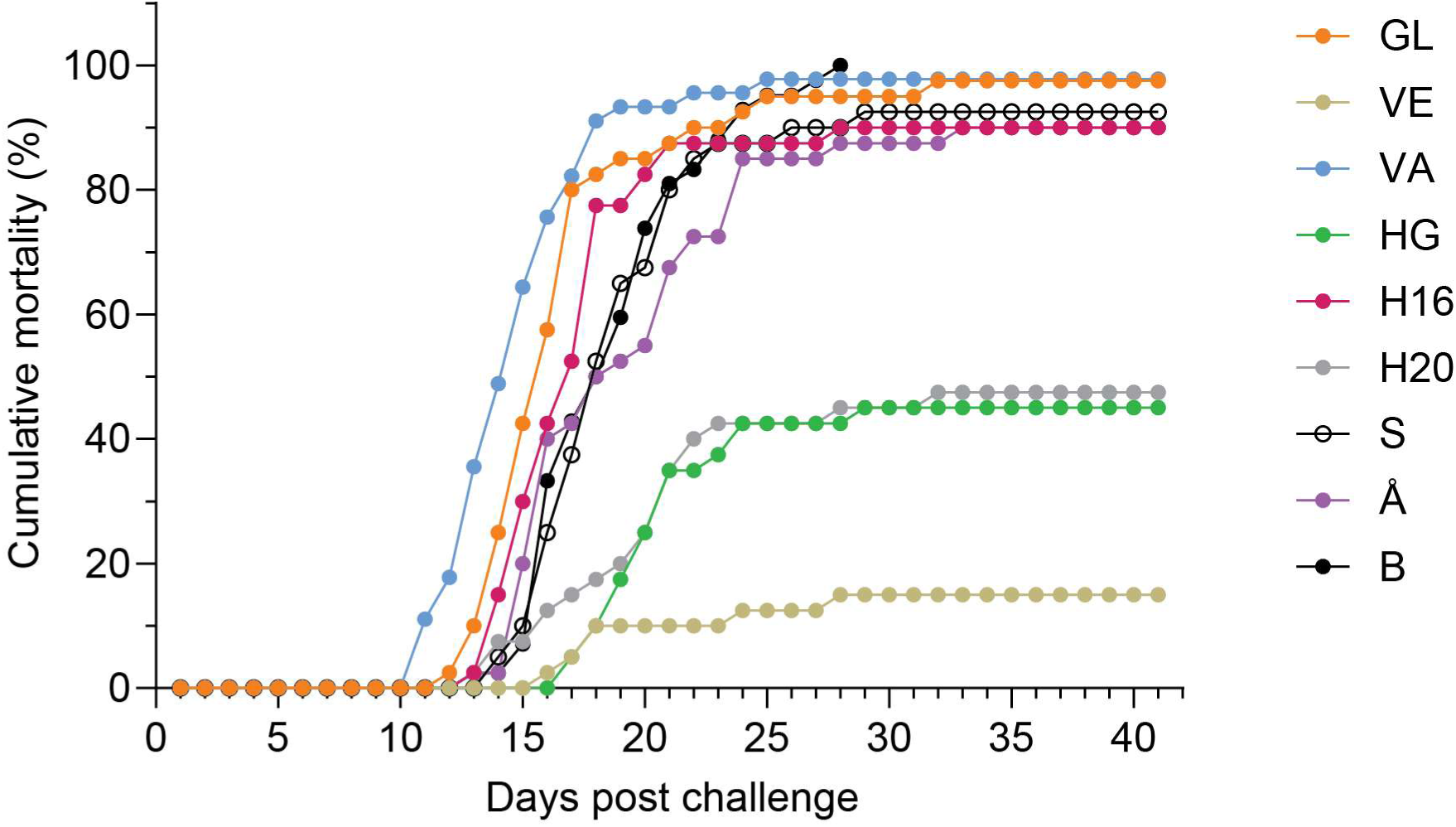
Overall cumulative mortality of Atlantic salmon in observation groups (n = 40 for GL, VE, HG, H16, H20, S, Å, n= 45 for VA, and n= 42 for B) following bath challenge with 10^3.5 TCID_50_/mL at 12 °C. Fish mortalities were observed for 42 days or until there were no more fish in the group. Note: Observation group was not included for L, thus mortality data for this isolate is not included.

Six of the nine isolates with OGs had mortalities above 90%. These isolates (GL, VA, H16, S, Å, and B) were classified as causing high mortality. Fish infected with HG and H20 showed mortality of 45% and 48% with onset of mortality at 13 and 17 DPC respectively. We designated these as causing moderate mortality. The last isolate VE with the first mortality on day 16 DPC and a cumulative mortality of 15% was designated as causing low mortality (Fig. 2). The L isolate could not be assigned to a mortality category; however, quantitative data of viral load in samples from L infected fish are shown together with isolates that caused moderate mortality. No mortalities were observed in the non-infected group in the trial period.

### Sequence variations among the ten ISAV isolates

Isolates in this study represented distribution in space and time over the last decade. Geographical spread along the coast of Norway for the ten isolates studied have been shown in Fig. 3.

**Figure 3.**
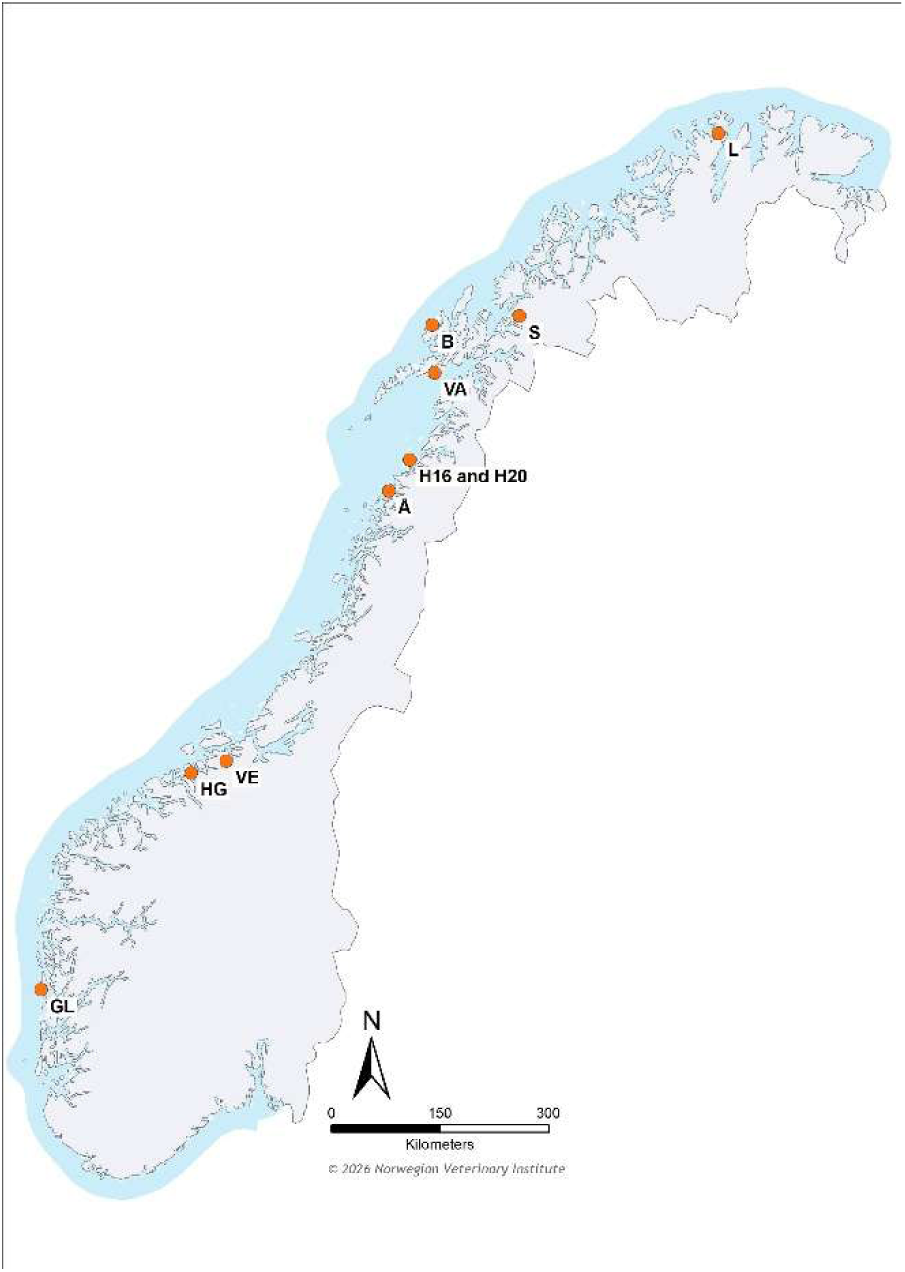
ISAV isolates chosen in this study showing geographical spread along the coast of Norway. Abbreviations have been described in Table 1. Map Illustration: Attilla Tarpai, Norwegian Veterinary Institute.

Amino acid sequences encompassing the haemagglutinin esterase HPR and the putative F protein cleavage site (residues ∼250–280) were aligned for the ten ISAV isolates, including the ISAV-HPR0 SK779 sequence as a reference for the full-length HPR. Sequence alignment (Fig. 4) revealed that eight of ten pathogenic isolates possessed the common pathogenicity associated with Q266L substitution and that VA contained an insertion of six residues (NNRFQRK) downstream of residue 266, both previously associated with virulence (40).

**Figure 4.**
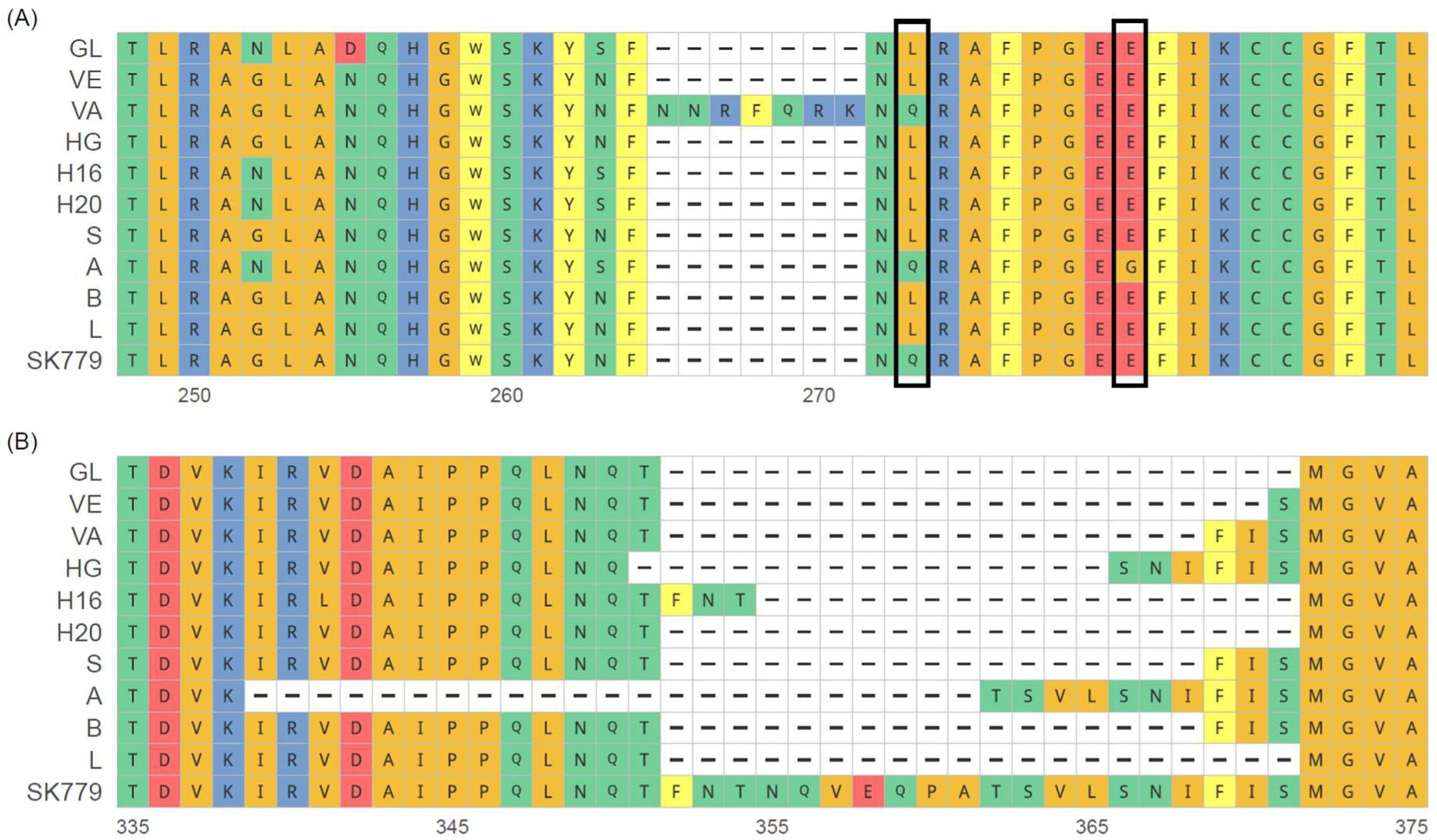
Multiple sequence alignment of the ISAV segment 5 region encoding the F protein cleavage site (A) and segment 6 region encoding the HE highly polymorphic region (B). The putative protease cleave sites is boxed in (A). The left box shows the canonical Q266L substitution while the right box shows the atypical E273G substitution. The coordinates up to the gaps are GL protein accession ADR77509.1 for (A) and ADR77510.1 for (B), respectively.

The severity of disease outbreaks is determined by the interaction between host, environment, and the infectious agent. Beyond the classical deletions in the HPR segment 6 and the Q266L mutation or insertions in the cleavage site on segment 5, we have limited or no knowledge of other mutations that contribute to virulence or differences in the virulence of the ILA virus in its ability to cause disease and disease severity. Isolate Å have a “HPR0-type” Q266 in combination with a E273G substitution within the conserved poly-glutamate region (Table 2) but still gave high mortality. This demonstrates that alternative point mutations close to the cleavage site can have comparable pathogenic effect to the Q266L point mutation or insertion. Our data confirms that pathotypic variants of segment 5 and segment 6 are required for invasive infection and the progression to ISA and widens the range of segment 5 variants that permit virulence. Moreover, the range of virulence in our trial cannot be explained by these changes alone, enforcing the commonly accepted notion that variation in other positions of the viral genome may contribute to ISAV virulence.

**Table 2.**
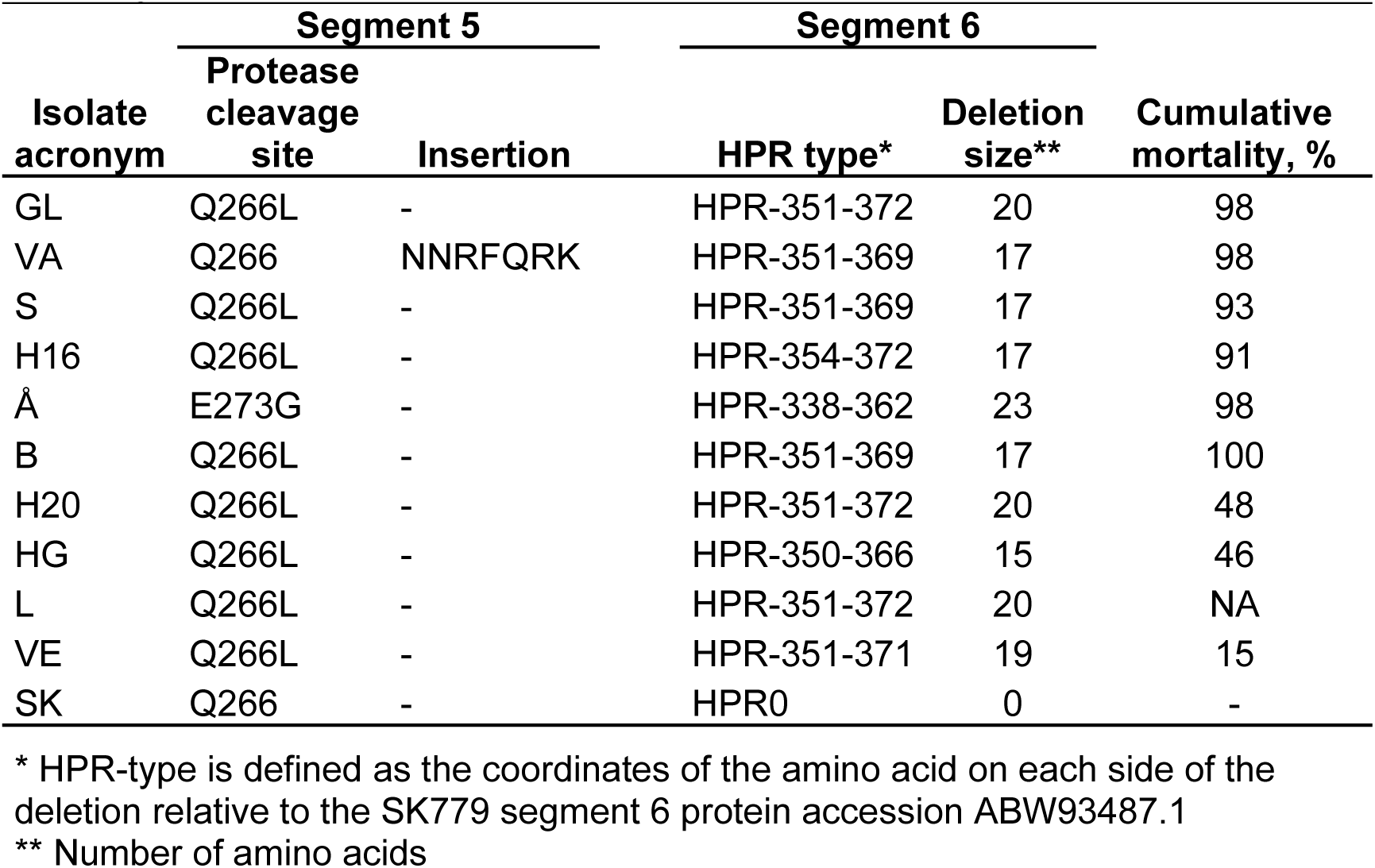
Properties of the putative protease cleavage site, the HPR and cumulative mortality of fish in their respective Observation Groups.

### Clinical Signs and Gross Pathology

The onset of clinical signs varied among the ten ISAV isolates. Upon observation, some fish appeared lethargic with reduced feeding activity. However, feed was found in the gut of most sampled individuals, suggesting normal appetite. External examination revealed the presence of haemorrhagic eyes, skin, and fins in some fish within each SG (Fig. 5A, B). Necropsy showed variable ascites in the peritoneal cavity of fish in most of the infected groups, with the onset of this observation varying from 7-28 DPC (Fig. 5A, C). Additional pathological findings included enlarged, friable liver, enlarged and dark spleen (Fig. 5C), swollen dark kidney and pale gills. No signs of disease or pathology were observed in the non-infected control group.

**Figure 5.**
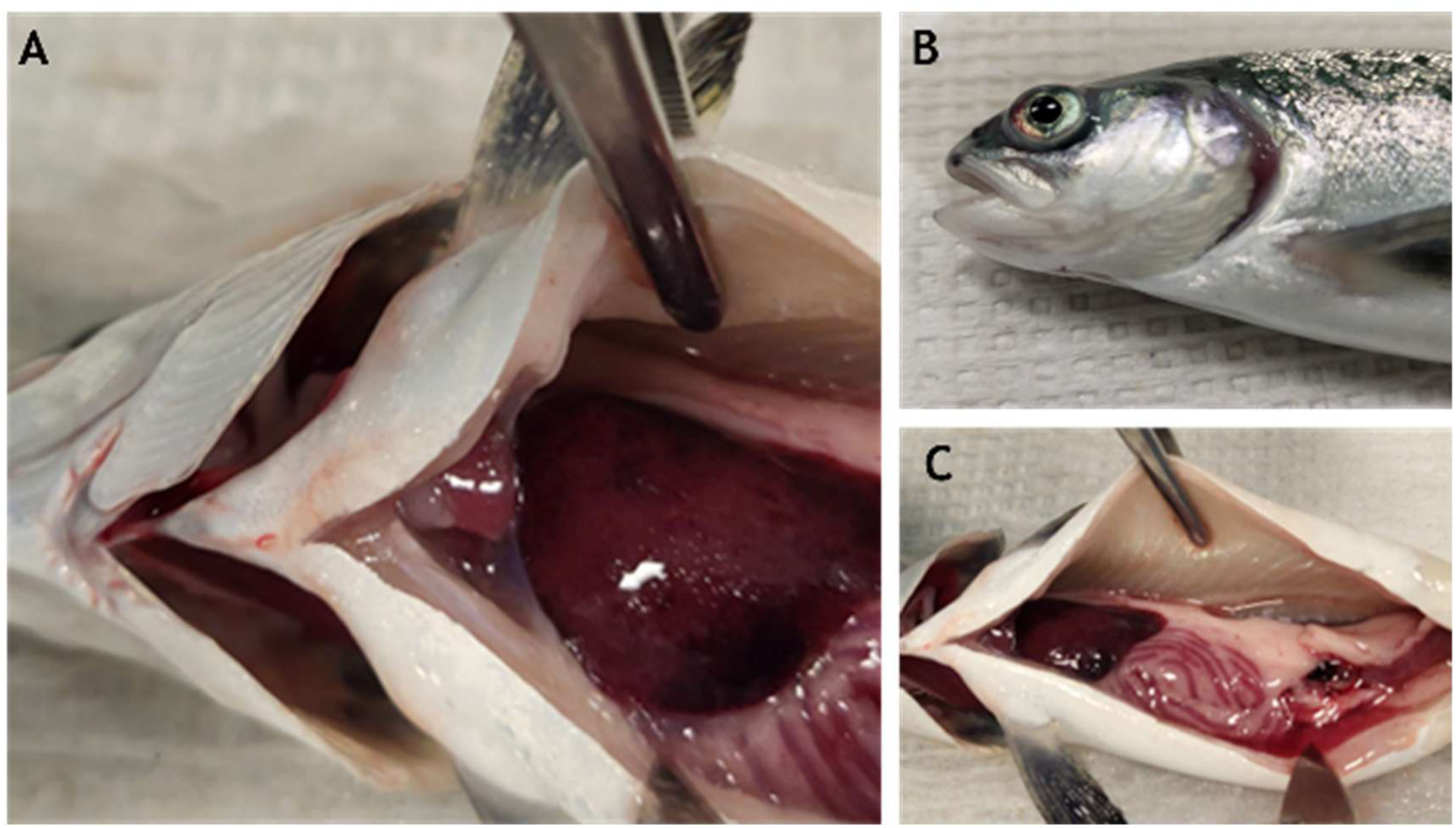
Necropsy showing classical ISA clinical signs in Atlantic salmon experimentally challenged with isolate GL as a representative for all isolates that gave high mortality. (A) gross pathology featuring enlarged, friable liver, and fluid in the heart sac (B) haemorrhagic eye, and (C) ascites, dark spleen and severe circulatory disturbances making the intestinal wall red.

### Quantification of ISAV RNA in Tank Water by RT-qPCR

Post challenge, tank water samples were collected at ten time-points, or until no fish remained in the SG tanks, to determine the presence of ISAV RNA in water by RT-qPCR. The levels of viral RNA in water differed markedly between isolates, in magnitude, timing of peak, and overall dynamics (Fig. 6). Viral RNA in water could be detected as early at 1 DPC for all isolates, with variable magnitude. Overall, the detection patterns fluctuated, with isolates peaking at different time points: 1 DPC for H16; 7 DPC for H20 ; 10 DPC for HG and VE; 14 DPC for VA; and 21 DPC for GL. Isolate H16 showed the highest and most rapidly detected levels of viral RNA in water with 6.7 × 10^5^ viral RNA cp per PCR reaction at 1 DPC, followed by Å (2.5 × 10^4^ ISAV RNA cp per PCR reaction) and H20 (1.4 × 10^4^ ISAV RNA cp per PCR reaction), respectively. The lowest detection at 1 DPC of 109 and 358 ISAV RNA cp per PCR reaction was detected for VA and S respectively. There was marked reduction in viral RNA in water at 3 DPC for several isolates, with continued decline at 7 DPC except for H20, H16, and Å, which showed a new peak at 7 DPC. Isolate S showed the lowest levels of viral RNA in water throughout the study period, peaking at 28 DPC (7.8 × 10^2^ ISAV RNA cp per PCR reaction). Isolate B showed high levels of viral RNA in water at 1 DPC but decreased markedly at 3 DPC, with minimal to no detection from 7 DPC onwards. Notably, the last fish was removed from the B SG at 21 DPC, precluding further sampling.

**Figure 6.**
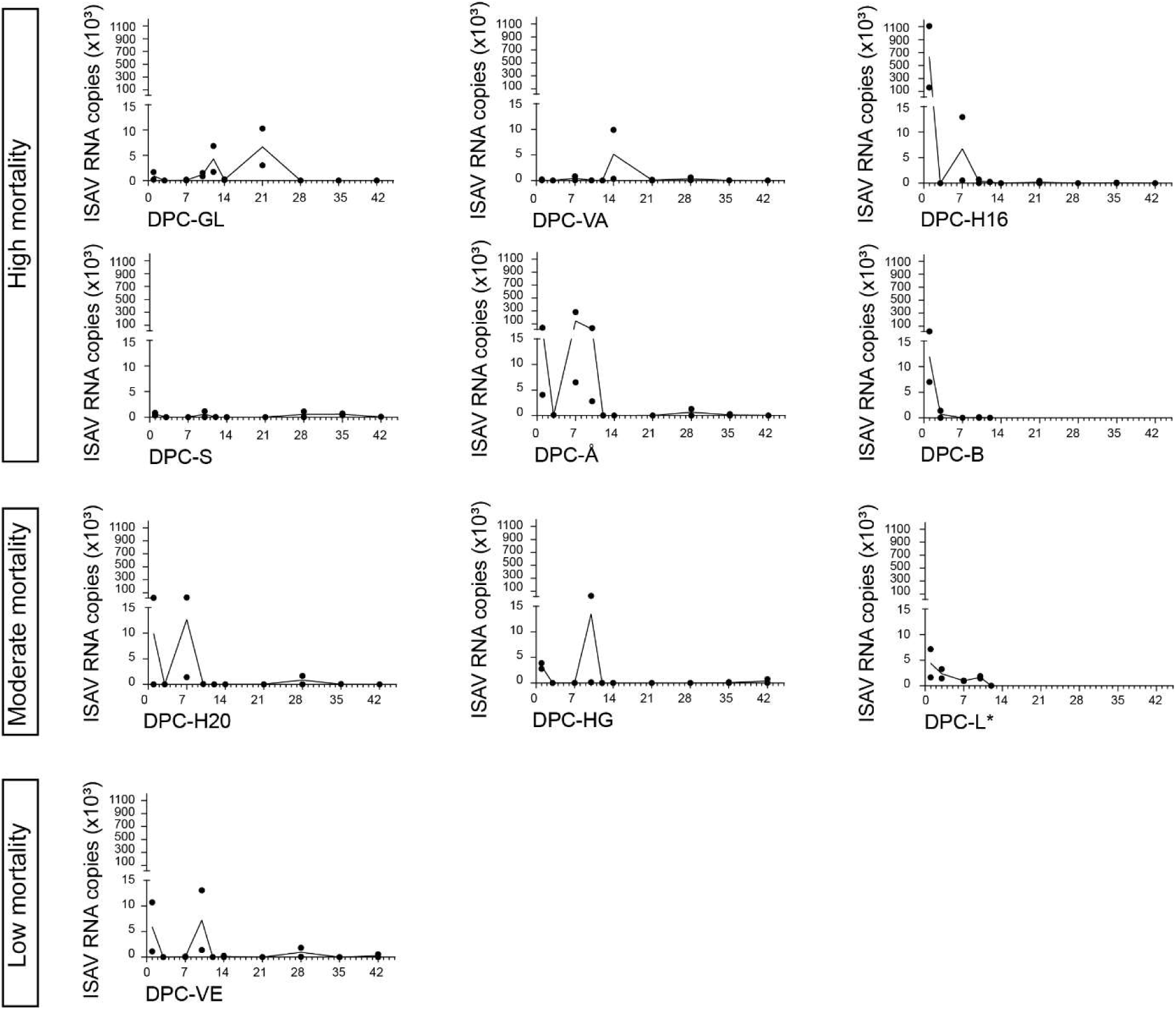
ISAV RNA in water samples for each isolate measured in SGs by RT-qPCR. N = 2 per time-point and copy numbers on Y-axis are represented per PCR reaction with 5 µL RNA. Isolate B reached 100% mortality at 28 DPC and no subsequent water analyses were performed for this isolate. The high mortality levels in some infection groups (B, GL, VA, S, H16, Å) combined with sampling from these tanks might have contributed to lower viral RNA levels observed for these isolates.

### Quantification of ISAV RNA load in fish gill tissue by RT-qPCR

ISAV RNA levels in gill tissue of challenged Atlantic salmon varied among isolates. Viral RNA was detected in gills for all isolates at 1 DPC, suggesting that viral attachment and initial replication occurred rapidly after exposure. For all isolates, the highest viral RNA loads at 1 DPC were detected in the gills, consistent with a local replication phase preceding systemic dissemination (41). Viral RNA levels had sharp increase in all groups from 3 to 7 DPC (Fig. 7).

**Figure 7.**
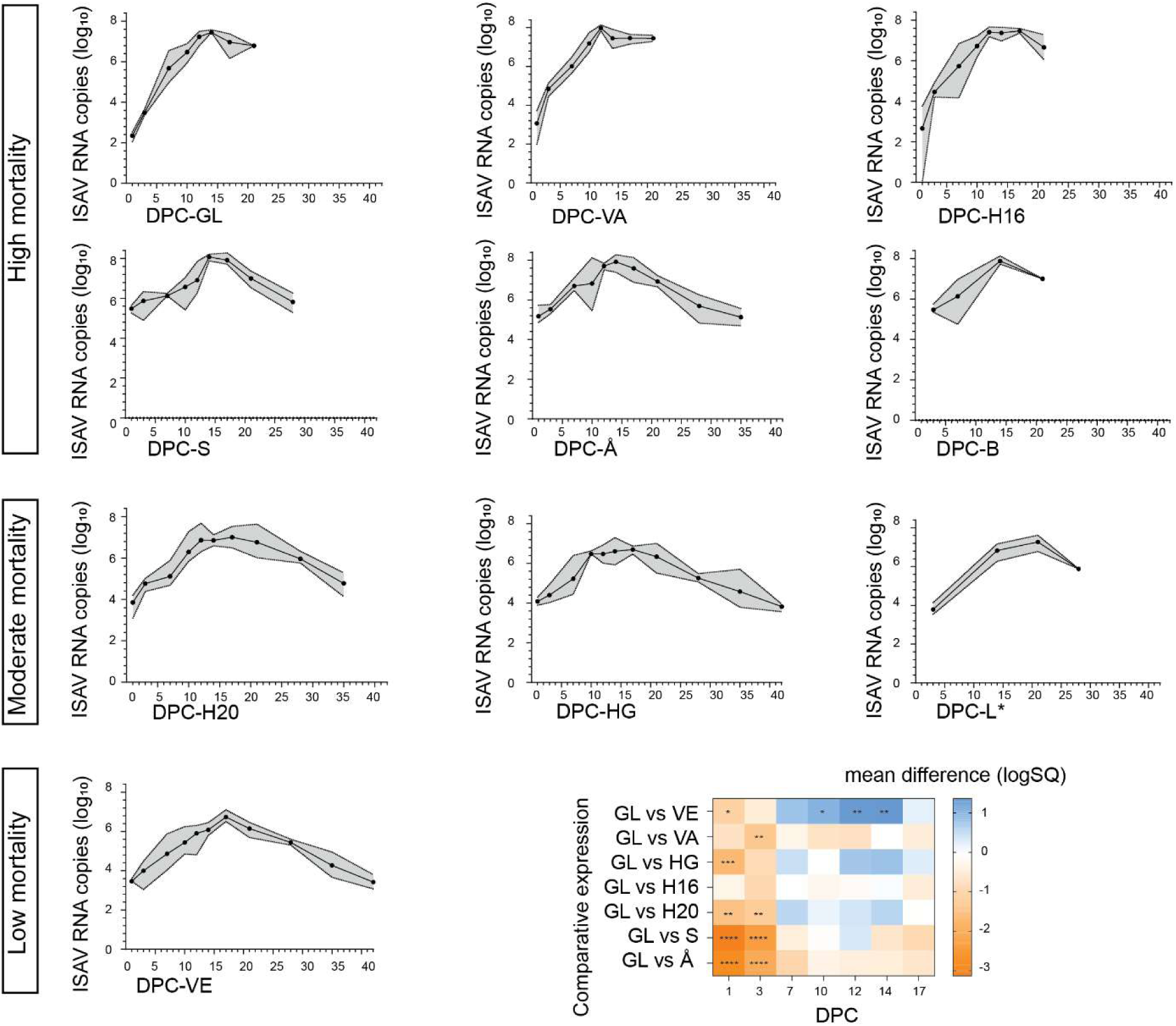
ISAV RNA load in the gill tissue samples for each isolate, measured by RT-qPCR. The graph for each isolate includes the period in which fish remained in the SG and could be sampled. N=4 per timepoint, except for GL, VA, and B at 21 DPC (N=2), S at 28 DPC (N=3), L at 28 DPC (N=2), and Å at 35 DPC (N=2). *Asterisk: Isolate L is grouped with moderate mortality isolates, but the lack of an OG precludes classification into a mortality category. Bottom right graph compares the results between fish groups infected with GL and each test isolate using two-way ANOVA with Dunnett’s multiple comparisons test. B and L were not included in the comparison, because of fewer sampling time points. *p<0.05, **p<0.01, ***p<0.001, ****p<0.0001.

The increase continued until 14 DPC for all groups apart from isolate VA where declining viral RNA was observed. Peak gill viral loads were generally observed between 12 and 14 DPC for isolates with high mortality, while the ones with moderate and low mortality peaked later, at 17 or 21 DPC (Table 3). Isolates with high mortality reached log_10_ 7.4–8.0 virus copies per PCR reaction, while for isolates H20, HG, L, and VE, highest viral RNA detected was log_10_ 6.7–7.3 virus copies per PCR reaction. All isolates maintained detectable viral RNA in gills until the last fish was removed from the SG tanks, which varied depending on mortality for each isolate (Fig. 7).

**Table 3.**
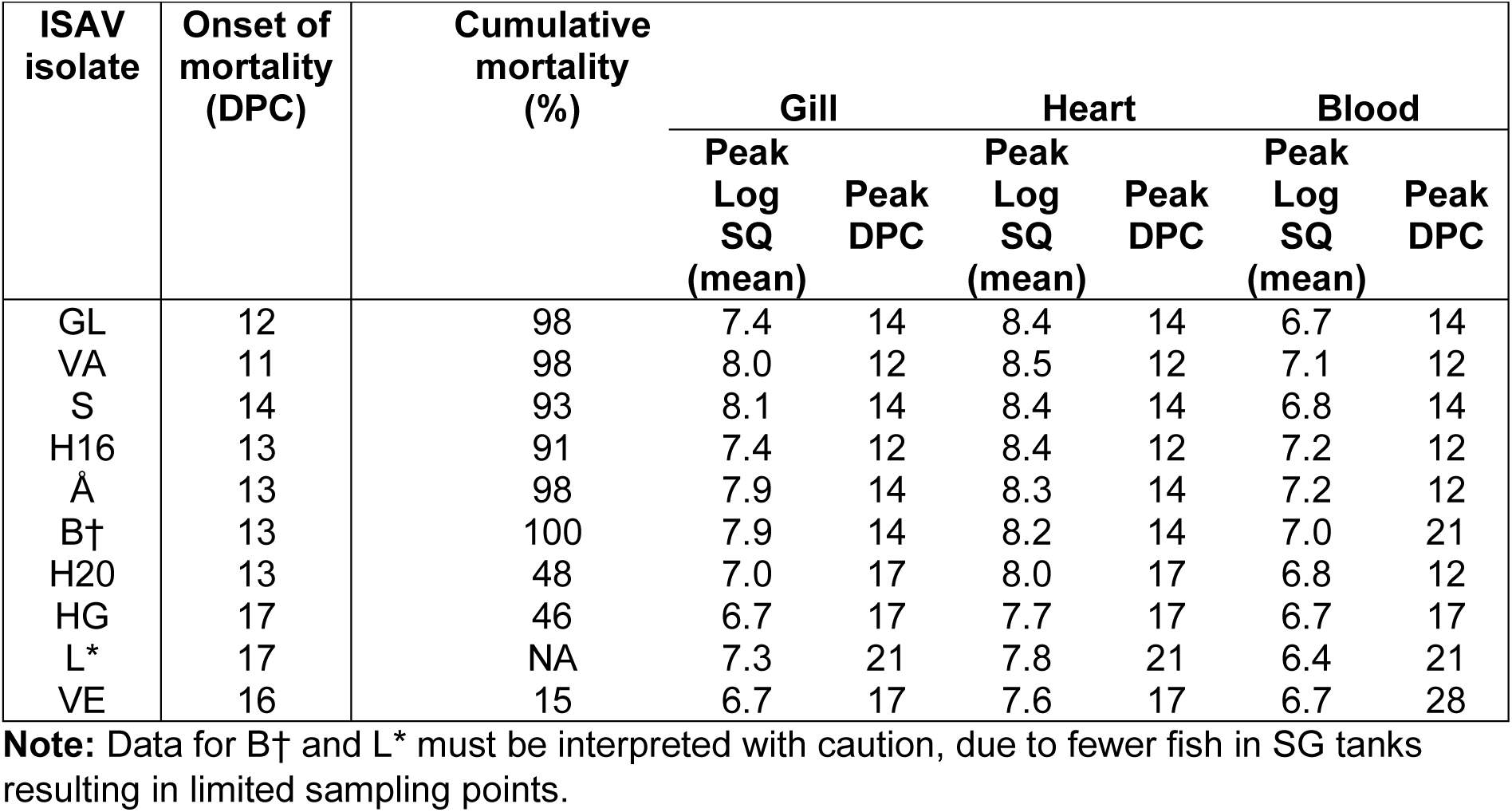
Comparative summary of gill, heart, and blood viral load kinetics, DPC: Days post challenge. For isolate H16 12 DPC value has been depicted in table, as it peaked 12 DPC with only slight increase to Log SQ 7.5 at 17 DPC.

| ISAV isolate | Onset of mortality (DPC) | Cumulative mortality (%) |  |  |  |  |  |  |
| --- | --- | --- | --- | --- | --- | --- | --- | --- |
|  |  |  | Gill |  | Heart |  | Blood |  |
|  |  |  | Peak Log SQ (mean) | Peak DPC | Peak Log SQ (mean) | Peak DPC | Peak Log SQ (mean) | Peak DPC |
| GL | 12 | 98 | 7.4 | 14 | 8.4 | 14 | 6.7 | 14 |
| VA | 11 | 98 | 8.0 | 12 | 8.5 | 12 | 7.1 | 12 |
| S | 14 | 93 | 8.1 | 14 | 8.4 | 14 | 6.8 | 14 |
| H16 | 13 | 91 | 7.4 | 12 | 8.4 | 12 | 7.2 | 12 |
| Å | 13 | 98 | 7.9 | 14 | 8.3 | 14 | 7.2 | 12 |
| B† | 13 | 100 | 7.9 | 14 | 8.2 | 14 | 7.0 | 21 |
| H20 | 13 | 48 | 7.0 | 17 | 8.0 | 17 | 6.8 | 12 |
| HG | 17 | 46 | 6.7 | 17 | 7.7 | 17 | 6.7 | 17 |
| L* | 17 | NA | 7.3 | 21 | 7.8 | 21 | 6.4 | 21 |
| VE | 16 | 15 | 6.7 | 17 | 7.6 | 17 | 6.7 | 28 |

**Note:** Data for B† and L\* must be interpreted with caution, due to fewer fish in SG tanks resulting in limited sampling points.

To compare ISAV RNA levels in gills across isolates we performed a two-way ANOVA analysis with Dunnett’s multiple comparisons between the type strain GL and the other isolates with equivalent sampling data (Fig. 7). Isolates B and L were excluded because of limited sampling. During the early phase of infection (1 and 3 DPC), gill viral loads were lower for GL than all other isolates, except H16. The greatest difference was observed at 1 DPC between GL and isolates S (mean difference log_10_-3.1, p<0.0001) and Å (mean difference log_10_ −2.8, p<0.0001). At later time points (7-17 DPC), VE was the only isolate that differed significantly from GL, showing reduced levels at 10 DPC (mean difference log_10_ 1.0, p<0.05), 12 DPC (mean difference log_10_ 1.3, p<0.01), and 14 DPC (mean difference log_10_ 1.4, p<0.01).

### Quantification of ISAV RNA load in heart tissues by RT-qPCR

Quantitative assessment of the replication kinetics of different isolates in the heart tissue showed that viral RNA levels varied among the isolates (Fig. 8). At 1–3 DPC, viral RNA was very low across most isolates (<10³ copies per PCR reaction, log_10_ < 3), reflecting limited systemic spread during the early phase of infection. GL was the only isolate with no detection of ISAV RNA in heart at 1 DPC.

**Figure 8.**
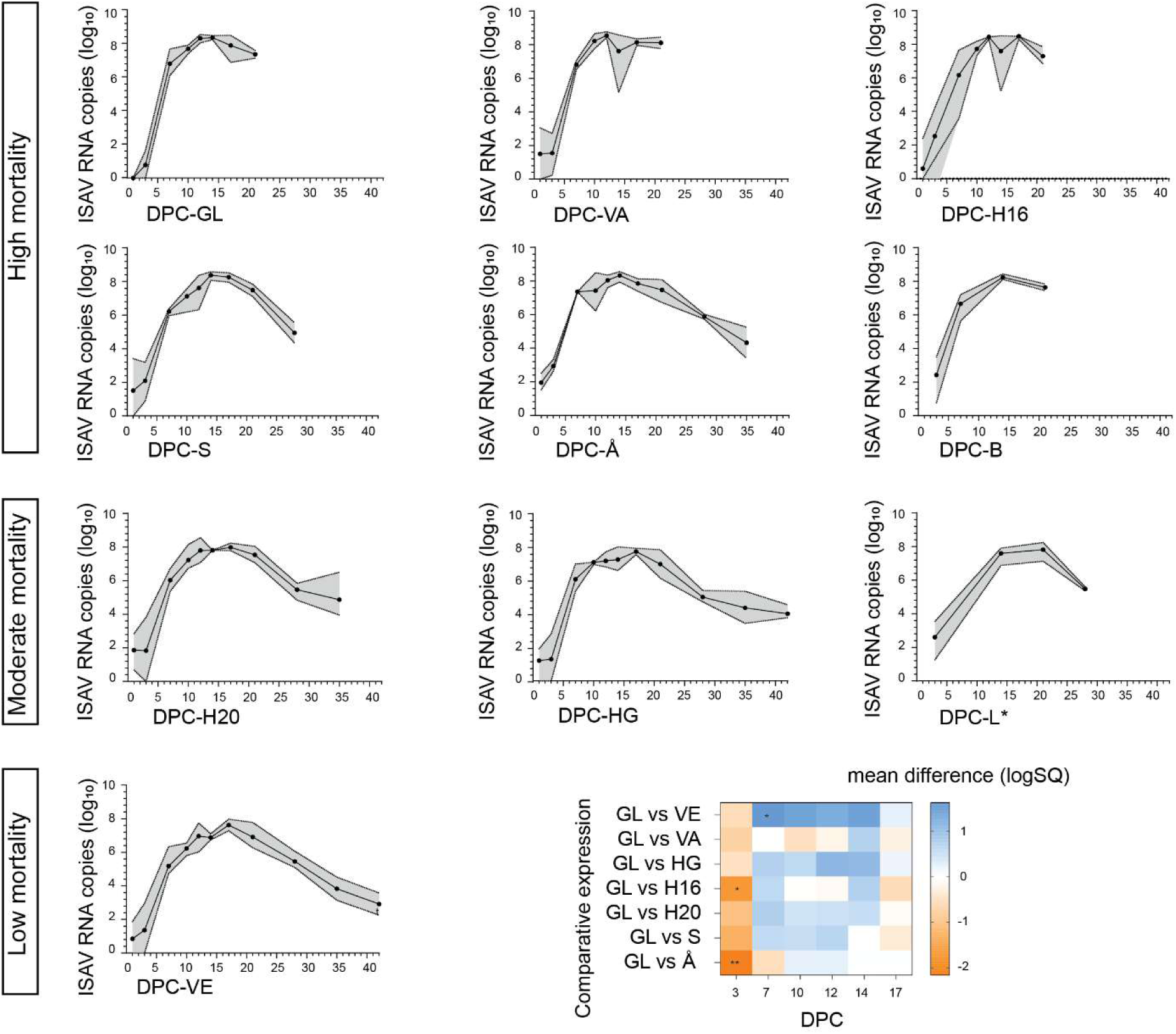
ISAV RNA load in the heart tissue samples for each isolate, measured by RT-qPCR. The graph for each isolate includes the period in which fish remained in the SG and could be sampled. N=4 per timepoint, except for GL, VA, and B 21 DPC (N=2), S 28 DPC (N=3), L 28 DPC (N=2), and Å 35 DPC (N=2). *Asterisk: Isolate L is grouped with moderate mortality isolates, but the lack of an OG precludes classification into a mortality category. Bottom right graph compares the results between fish groups infected with GL and each test isolate using two-way ANOVA with Dunnett’s multiple comparisons test. B and L were not included in the comparison, because of fewer sampling time points. *p<0.05, **p<0.01.

The viral load in heart increased exponentially for all isolates until 14 DPC. All isolates with high mortality reached a peak ranging between log_10_ 8.2 to 8.5 viral RNA copies per PCR reaction by 14 DPC (Table 3). Peak log₁₀ value for H20, HG, L, and VE, were 7.6 – 8.0 by 17 - 21 DPC. Peak heart viral loads were generally observed between 12 and 14 DPC for isolates with high mortality, while the ones with moderate and low mortality peaked later, at 17 or 21 DPC (Table 3). After 21 DPC, viral loads gradually declined. Viral RNA in heart remained detectable until the termination of experiment at 35 or 42 DPC or until the last fish was removed from SG tanks.

To compare ISAV RNA levels in heart across isolates we performed a two-way ANOVA analysis with Dunnett’s multiple comparisons between the type strain GL and the other isolates with equivalent sampling data (Fig. 8), excluding data from 1 DPC when no virus RNA was detected in GL. Isolates B and L were excluded because of limited sampling. During the early phase of infection (3 DPC), heart viral loads were lower for GL than H16 (mean difference log_10_ −1.8, p<0.05) and Å (mean difference log_10_ −2.2, p<0.01). At 7 DPC, VE was the only isolate that differed significantly from GL, showing reduced levels (mean difference log_10_ 1.6, p<0.05). Between 10 and 17 DPC no isolate showed a significant difference in ISAV RNA levels in heart tissues to GL, suggesting overall convergence over time.

### Quantification of ISAV RNA load in whole blood by RT-qPCR

ISAV RNA was not detected in blood at 1 DPC of any of the virus isolates, except for H16. By 3 DPC, low levels of ISAV RNA (<log 2) were detected in all isolates, except for VE, HG and H20, marking the onset of systemic spread. All isolates exhibited a pronounced rise in viral RNA in blood between 3 and 14 DPC (Fig. 9). Isolates GL, S, H20, HG and VE reached peak log₁₀ values of 6.7–6.8, H16, B, VA, and Å peaked between 7.0–7.2, while L showed lowest levels at log 6.4 (Table 3). For most isolates, ISAV RNA detected in blood reached its maximum around 12–14 DPC, followed by a plateau or slow decline at 14–21 DPC. Isolate VE showed very delayed peak of viral RNA in blood at 35 DPC. Time-point for decrease in viral RNA varied between the isolates, and maintaining detectable levels up to the end of trial or alternatively until last surviving fish in the SG tanks were sampled suggesting prolonged circulation (Fig. 9).

**Figure 9.**
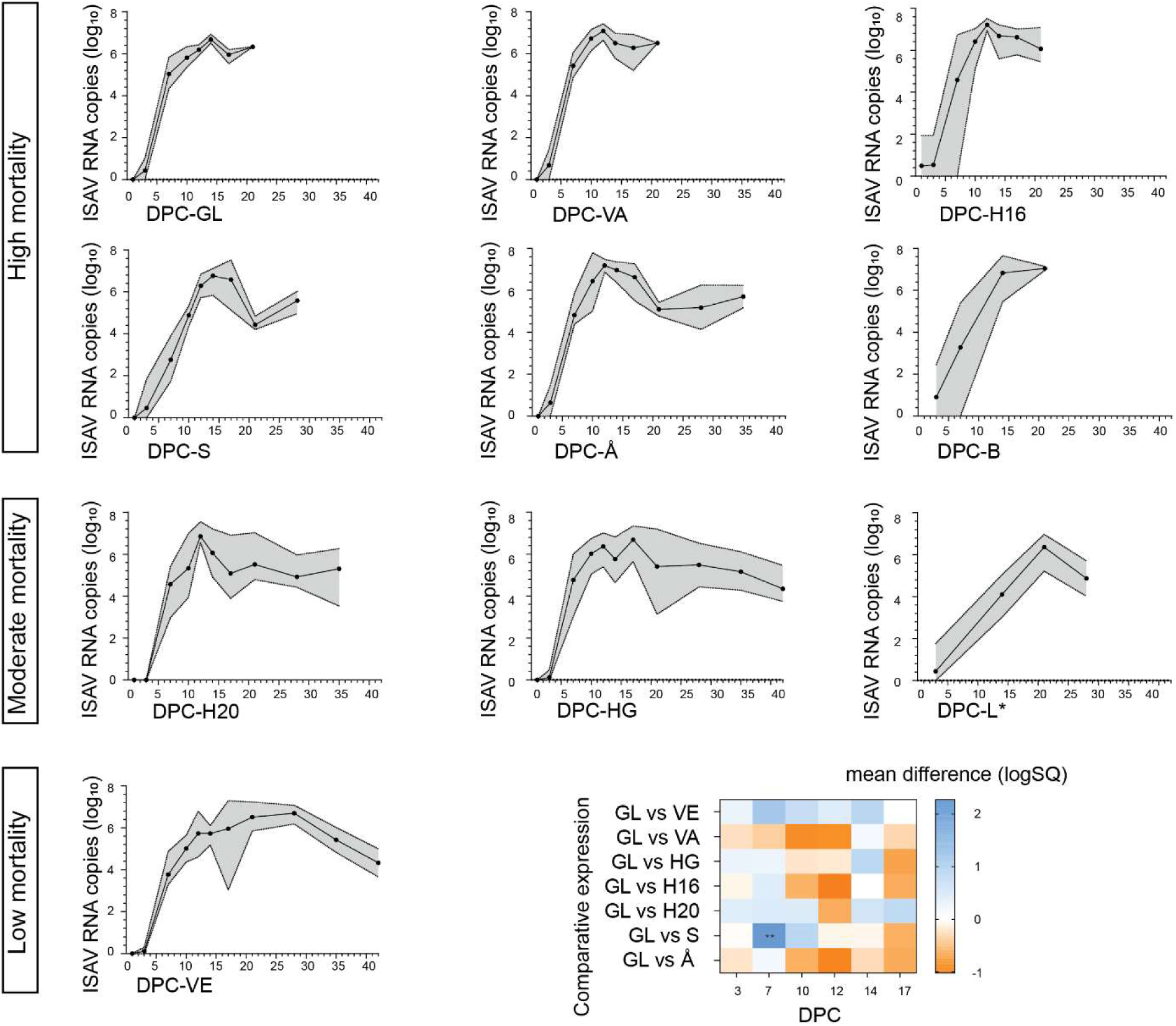
ISAV RNA load in the blood samples for each isolate, measured by RT-qPCR. The graph for each isolate includes the period in which fish remained in the SG and could be sampled. N=4 per timepoint, except for GL, VA, and B 21 DPC (N=2), S 28 DPC (N=3), L 28 DPC (N=2), and Å 35 DPC (N=2). *Asterisk: Isolate L is grouped with moderate mortality isolates, but the lack of an OG preludes classification into a mortality category. Bottom right graph compares the results between fish groups infected with GL and each test isolate using two-way ANOVA with Dunnett’s multiple comparisons test. B and L were not included in the comparison, because of fewer sampling time points. **p<0.01.

To compare ISAV RNA levels in blood across isolates we performed a two-way ANOVA analysis with Dunnett’s multiple comparisons between the type strain GL and the other isolates with equivalent sampling data (Fig. 9), excluding data from 1 DPC when no virus RNA was detected in GL. Isolates B and L were excluded because of limited sampling. The only isolate that showed a statistically significant difference in ISAV RNA levels in blood compared to GL was S, with reduced levels at 7 DPC (mean difference log10 2.3, p<0.01). For remaining isolates and time points, ISAV RNA levels in blood were comparable to GL.

### Flow-cytometric characterization of the dynamic binding of ISAV to blood cells

During viraemia, erythrocytes (RBC) bind ISAV particles and occasionally express ISAV HE (42). While other blood cell types potentially could bind ISAV, they make up a minor proportion of circulating cells and are unlikely to contribute to the results of a flow cytometric analysis, which gives comparable results to quantifying ISAV-bound RBC by manual counting (39). The percentage of blood cells carrying detectable levels of ISAV particles and/or expressing surface ISAV HE in infected fish was determined by flow cytometry. The amplitude of fluorescent signal was not directly comparable across isolates, as we could not exclude potential strain-specific differences in antibody affinity, but allowed comparison of the binding dynamics. The initial ISAV-RBC binding dynamics mostly reflected the levels of viral RNA in the blood, with VE showing a delayed onset of binding (Fig. 10). A similar delay was observed for Å, despite the early detection of this isolate in the blood by RT-qPCR. For all isolates, the degree of ISAV binding fell sharply over time, again with Å showing a more rapid decrease than the other isolates. Very limited ISAV binding to cells was observed after 28 DPC, even for VE that showed the highest levels of viral RNA in blood at 28-35 DPC.

**Figure 10.**
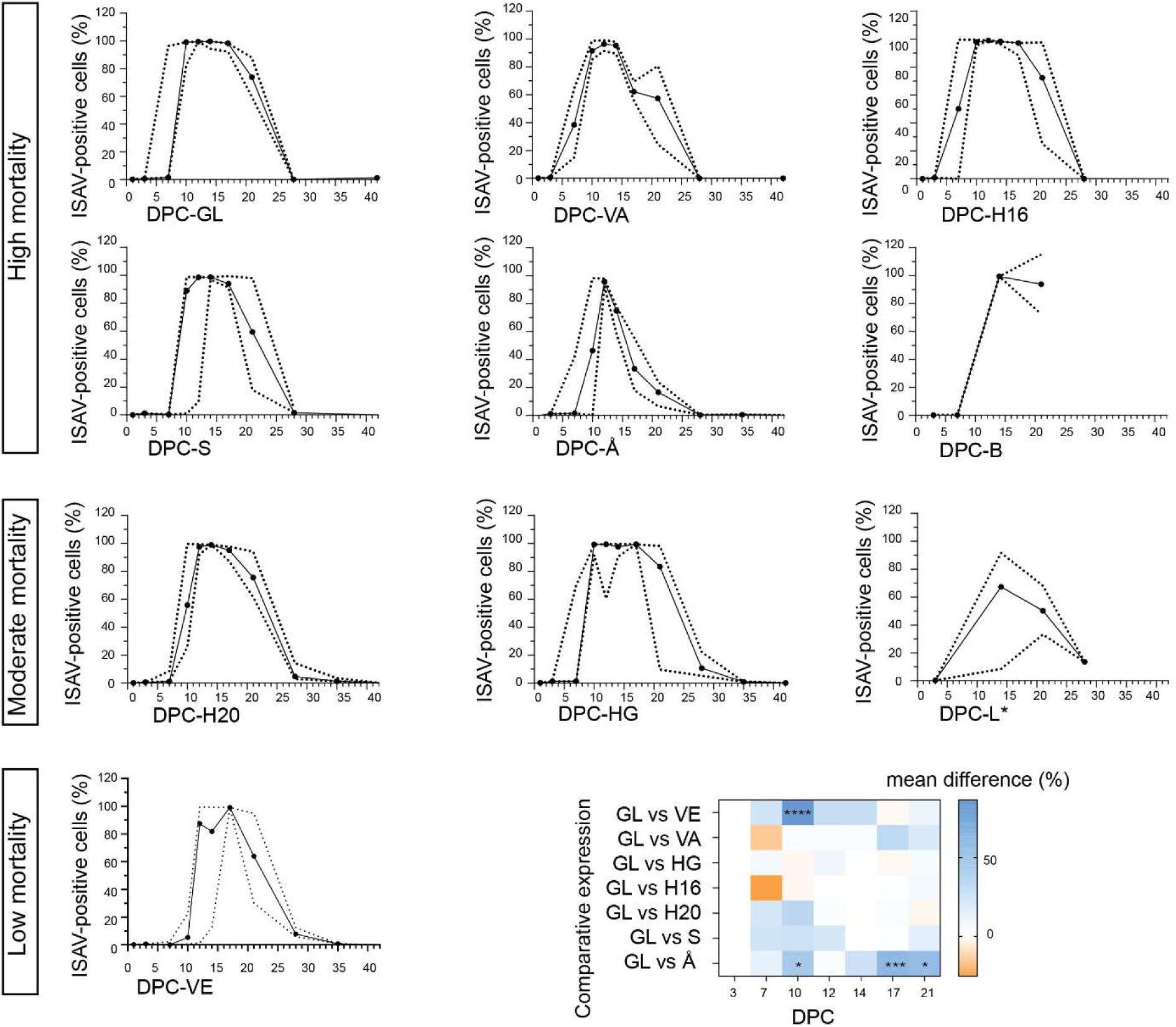
Quantification of ISAV-positive blood cells, measured by flow cytometry. Bottom right graph compares the results between fish groups infected with GL and each test isolate using two-way ANOVA with Dunnett’s multiple comparisons test. B and L were not included in the comparison, because of fewer sampling time points. *p<0.05, ***p<0.001, ****p<0.0001. N = 4 per time-point, except GL, VA, and B 21 DPC (N=2), S 28 DPC (N=3), L 28 DPC (N=2), Å 35 DPC (N=2), GL and VA 42 DPC (N=1), VE 42 DPC (N=21), HG 42 DPC (N=14), H20 42 DPC (N=6), and S 42 DPC (N=3). *Asterisk: Isolate L is grouped with moderate mortality isolates, but the lack of an OG preludes classification into a mortality category.

To compare the percentage of ISAV-positive blood cells across isolates we performed a two-way ANOVA analysis with Dunnett’s multiple comparisons between the type strain GL and the other isolates with equivalent sampling data (Fig. 10), excluding data from 1 DPC when no virus RNA was detected in GL. Isolates B and L were excluded because of limited sampling. No significant difference between GL and the other isolates was observed before 10 DPC, when the percentage of ISAV-positive blood cells was higher in GL than VE (mean difference 87%, p<0.0001) and Å (mean difference 47%, p<0.05).

At 12 and 14 DPC, the percentage of ISAV-positive RBC was comparable to GL for all isolates, while Å showed reduced levels at 17 DPC (mean difference 62%, p<0.001).

### Histopathology and Immunohistochemistry

Histopathology showed that all the ISAV isolates caused pathology in accordance with previous descriptions of ISA, including focal cellular changes in the spleen (Fig. 11A), characterized by localized alterations in splenic tissue architecture; circulatory disturbances and signs of RBC breakdown in the kidney (Fig. 11B); and haemorrhagic liver necrosis (Fig. 11C). Evidence of haemorrhagic liver necrosis, considered a near pathognomonic lesion of ISA, was found in all isolate groups apart from the low mortality isolate VE.

**Figure 11.**
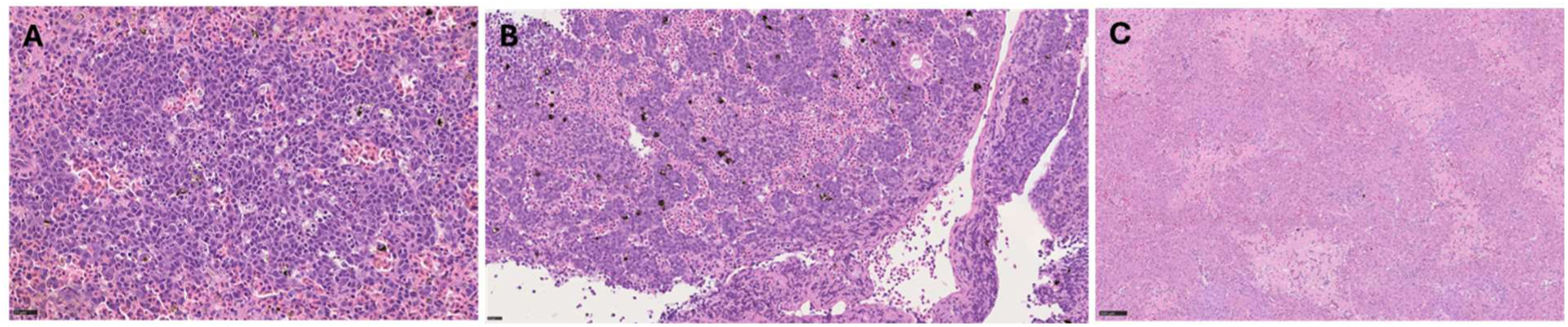
Pronounced histological lesions in H&E-stained tissues from GL-infected Atlantic salmon. (A) Spleen with focal cellular change, 14 DPC (B), Kidney with circulatory disturbances,12 DPC, and (C) liver with zonal necrosis and focal haemorrhage, 21 DPC. Scale bars A-B = 25µm and C = 100µM.

However, one VE-infected fish sampled at 28 DPC had sparse focal liver necrosis as well as clear signs of increased RBC breakdown, indicating that this isolate can cause ISA related pathology. The sum of average pathology scores presented in Table 4 align well with mortality, showing that high mortality corresponds to more severe pathology in the period 7-28 DPC.

**Table 4.**
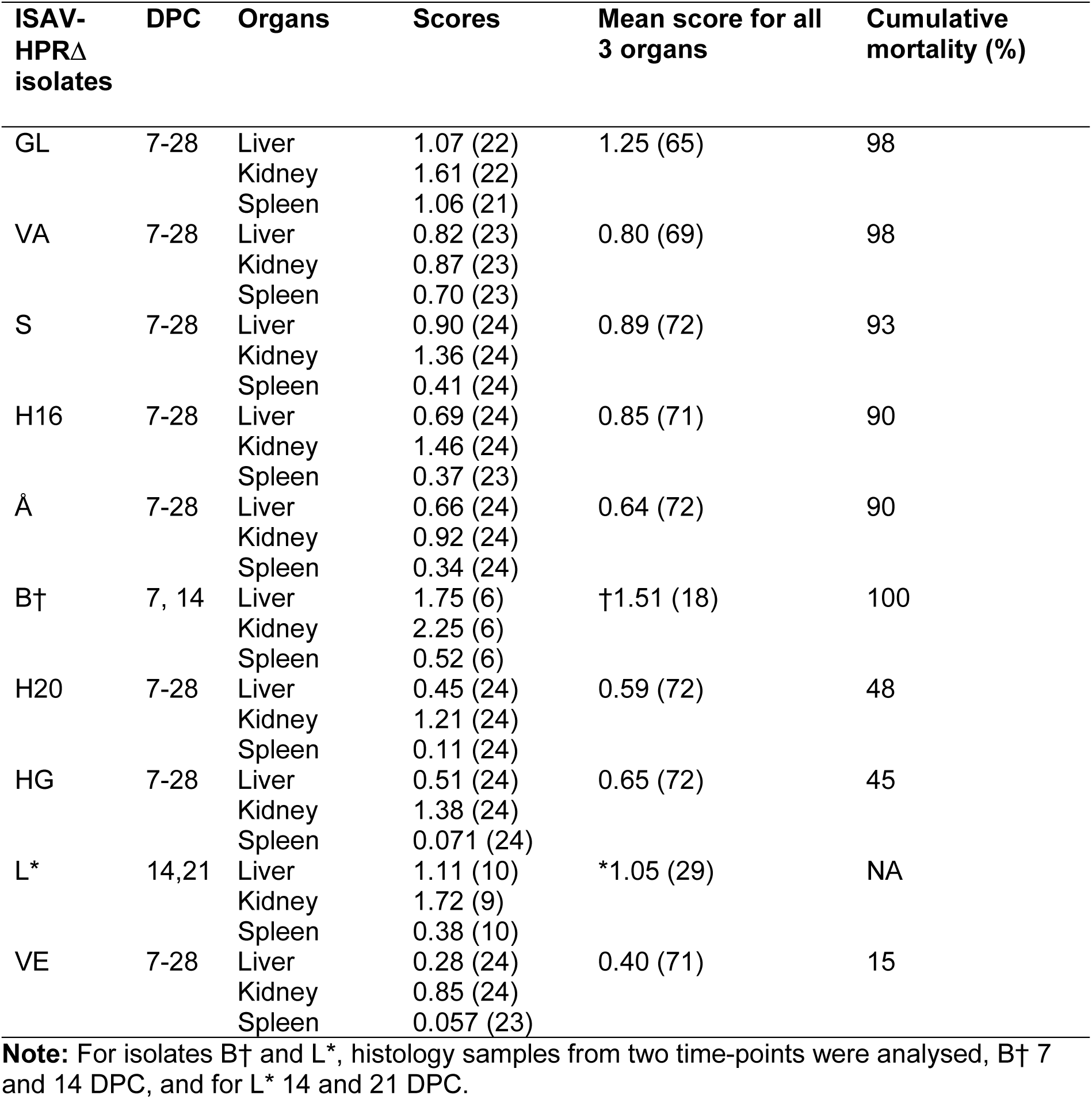
Histopathology scores of main target organs of fish infected with each ISAV isolate. Numbers in parentheses represent total fish analysed. For B and L – number of fish in Sampling group (SGs) was 24 and 21 respectively. NA – Not applicable, as Observation Group (OG) was not included for L, thus mortality data for this isolate is not included.

The IHC of heart, and spleen tissue samples from two fish per sampling time point from each SG showed ISA antigen distribution according to the endotheliotropic nature of ISAV-HPRΔ (Fig. 12). The vascular endothelium in the heart was most consistently and severely infected. Within the heart tissue, the valves and the bulbus appeared to be predilection sites of infection, although infection with e.g. GL appeared to involve the entire heart endothelium at the peak of infection (Fig. 12).

**Figure 12.**
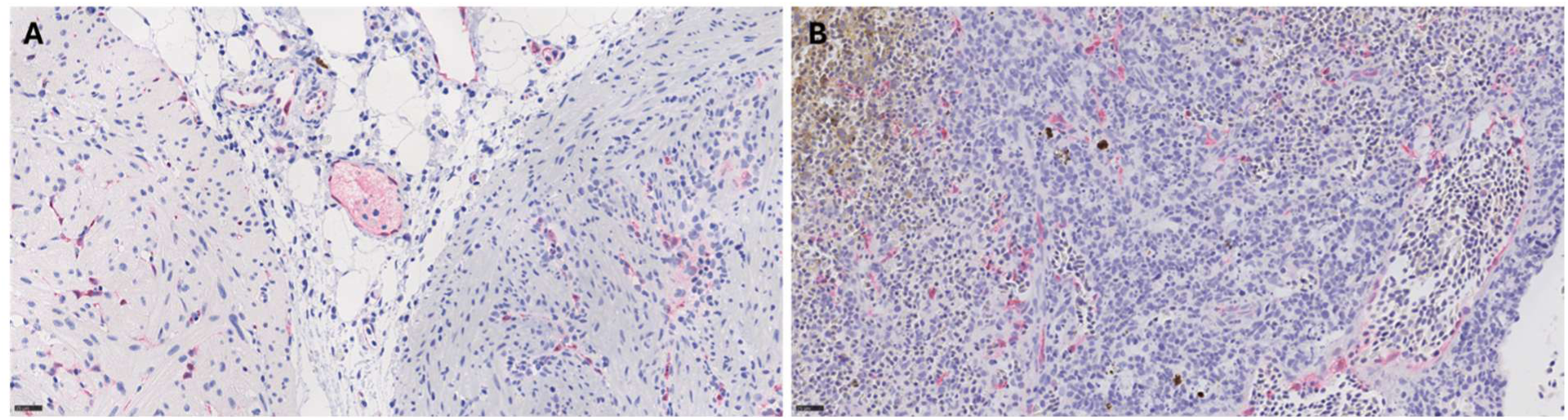
Immunohistochemical staining of (A) heart and (B) spleen from GL-infected Atlantic salmon at 21 DPC, showing widespread endothelial distribution of ISAV NP in (red) (A) heart, with the compact muscular layer to the left and bulbus to the right and (B) spleen. Scale bars A-B = 25µm.

Both isolate Å, giving high mortality, and H16 and HG, giving moderate mortality, showed a comparable pattern of histopathological lesions and IHC staining, but less pronounced than the other high mortality isolates. Fish infected with VE primarily showed signs of increased and prolonged RBC breakdown with sparse ISAV staining in endothelium. No signs of ISA-typical pathology or ISAV antigen were observed in noninfected control fish.

## Discussion

This study provides a comprehensive comparative analysis of infection kinetics, tissue tropism, virus shedding, pathology, and mortality caused by ten ISAV-HPRΔ isolates, including the well-characterized highly virulent Norwegian reference strain NO/Glesvaer/2/90 (GL) and nine recent Norwegian field isolates. Using a standardized bath challenge model, our findings reveal substantial biological variability among ISAV-HPRΔ isolates, suggesting that ISAV-HPRΔ isolates exhibit a wide spectrum of infection dynamics and mortality. Mortality, onset of mortality, and shedding did not correlate, suggesting that these outcomes, that are important for viral virulence, may be determined by distinct viral properties.

### Variation in virulence and mortality patterns

The ISAV isolates caused a broad spectrum of mortality outcomes, ranging from low (15%) to very high (100%). The time of the first mortality events varied between the isolates and occurred between 11 – 17 DPC, consistent with previous studies (14, 36). All isolates apart from HG and VE induced rapid onset of mortality between 11 and 14 DPC, however H20 did not cause high cumulative mortality despite early onset of mortality. Isolate HG and VE displayed delayed onset resulting in moderate cumulative mortality for HG and low mortality of 15% for VE. This indicates a lack of correlation between the initial time of onset of mortality and cumulative mortality at the isolate level. It is difficult to align these outcomes with field observations as mortalities in the field can be influenced by numerous environmental and host factors. In addition, to limit the spread of ISAV, fish populations with confirmation of ISA are often culled before mortality reach high levels. Reliable estimation of mortality across isolates therefore requires the use of a standardized challenge model. Our study shows that five out of eight recent field isolates (B, VA, H16, S, and Å) caused comparative morbidity and mortality to the type strain GL. The trial also shows that not all ISAV-HPRΔ infections result in severe mortality, despite their common genetic classification as “pathogenic” by WOAH (43). This diversity reflects that observed in a previous study comparing ISAV isolates from the 1990s (44). Overall, our results suggest that the pathogenic potential of currently circulating ISAV strains is comparable to the strains at the peak of the Norwegian ISA epidemic circulating in 1990s.

As for all pathogenic orthomyxoviruses, the observed differences in mortality likely result from a combination of several factors, including variations in viral replication efficiency in internal organs, systemic dissemination rates, environmental conditions, stress, and immune evasion. High and moderate mortality isolates produced rapid systemic dissemination. In contrast, isolate VE resulted in lower viral loads, limited tissue involvement and relatively low mortality, despite pathotypic changes in segment 5 and 6. Detection of VE at the farm was not associated with clinical signs of ISA, suggesting that its low mortality was, at least in part, attributable to intrinsic viral factors. The two isolates that caused moderate mortality in our infection trial, HG (46%) and H20 (48%), were likewise isolated from farms reporting low mortality.

The virus isolates exhibited genetic diversity, including the pathotypic changes in segment 5 and segment 6 (40). Beyond the classical deletions in the HE HPR and QL266 or insertions in the F protein, we have limited or no knowledge of mutations that contribute to differential virulence in ISA.

Our data indicate that a wider range of alterations in the genomic region encoding the F protein cleavage site in addition to a deletion in the HE gene may be sufficient for ISAV virulence. Both Q266L mutations and insertions, like the insertion detected in the highly virulent (98%) VA isolate (NNRFQRK), are widespread and may enhance pathogenicity, potentially by facilitating more efficient proteolytic activation of the F protein. Notably, isolate Å featured no Q266L substitution or insertion but a unique E273G substitution and still gave high mortality. This demonstrates that alternative substitutions near the F protein cleavage site can mediate pathogenicity, highlighting the need to understand the molecular mechanism by which changes in this region influence infection. Our data confirms that pathotypic variants of segment 5 and segment 6 are required for invasive infection and the progression to ISA and widens the range of segment 5 variants that permit virulence. Moreover, the range of virulence in our trial cannot be explained by these changes alone, enforcing the commonly accepted notion that variation in other positions of the viral genome may contribute to ISAV virulence (1, 45). Notably do GL, H20 and L have identical HE deletion and F substitution while showing clear differences in mortality.

### Virus shedding dynamics and transmission potential

Quantitative PCR analysis of tank water has provided novel insights into the dynamics of virus shedding. Although it is possible to detect infective ISAV particles in water samples (46), testing all water samples from all ten tanks for infective particles at each sampling time point was not practically feasible in the current study. While the amount of viral RNA in water may not reflect the presence of infective virus particles, it still provides a reasonable proxy to estimate transmission potential. The differences in average viral RNA load in water among isolates were striking. Isolate H16 demonstrated high and rapid detectable viral RNA in water, indicating potentially efficient replication and environmental release, whereas isolates S, L, and VA showed minimal shedding, despite efficient replication in epithelium. High shedding may correlate with enhanced transmission risk, particularly under farm conditions with shared water systems. The detection of viral RNA in tank water as early as 1 DPC for all isolates may suggest rapid virus-host contact and initial shedding. The levels of viral RNA were higher in gills than heart and blood for all isolates at early time points (1-3 DPC), supporting the hypothesis of local propagation in gills and shedding during early infection. The low replication in gill observed for GL at 1 DPC aligns with previous observations (47, 48). Caution should be applied when interpreting these data. While it is possible that the viral RNA detected in water 1 DPC represents rapid virus propagation in gills, it needs further substantiation. Since all fish were bath infected in one tank and then transferred to their designated tanks, detection attributed to remaining virus that was added for bath infection can be ruled out. However, it can be speculated that virus adhering to fish mucosal surfaces during infection could have been released back into the water after transfer. We observed considerable differences in average viral RNA load in water among isolates. Isolate H16 demonstrated high and rapid detectable viral RNA in water, potentially indicating efficient replication and environmental release, whereas isolates S, L, and VA showed minimal shedding throughout the trial, despite efficient replication in gill epithelium. The shedding profiles did not seem to be closely related to the onset of mortality, percentage of cumulative mortality or viral load in target organs. Isolate B with highest mortality had moderate shedding at 1 DPC but drastically reduced at later timepoints. Isolates VA and GL, with high mortality, showed limited and moderate shedding, while H16 and Å reached highest viral loads but lower mortality than B, GL and VA.

Detection of viral RNA in water from early phase of infection, before signs of disease or mortality develop, until late phase suggests that one might increase the risk of transmission if intervention depends on confirmation of ISA in a population. Variation in ISAV RNA in water over time and between isolates observed in our study suggests that environmental nucleic acids (eNA) in water should be carefully assessed when applying for risk of transmission or for ISAV surveillance. We need to carefully assess and consider several factors. In our study we measured virus shedding in SGs where the number of fish declined over time not only due to mortality, but also due to serial sampling. This might have influenced analysis of shedding rate and amounts, and thus systematic surveillance should be carried out to understand the applicability of eNA surveillance. Our results show lack of correlation between mortality and ISAV RNA in water, though there was a positive correlation with gill viral RNA especially at early time-points (1-3 DPC). This is interesting as it has been established earlier that ISAV-HPR0, which mainly propagates in gills, seems to be highly efficient at transmission, but lacks virulent potential (49).

### Tissue pathology

As expected, histopathological and immunohistochemical analyses confirmed the endotheliotropic nature of ISAV-HPRΔ infection. The lesions observed were consistent with classical ISA pathology, suggesting circulatory disturbances (50). Lesion severity and tissue distribution varied between isolates, but the severity of histopathological findings largely mirrored the cumulative mortality. The IHC findings further supported these observations with detection of ISAV antigen in vascular endothelium with the intensity indicating higher viral loads. Interestingly, the moderate mortality isolates (H20 and HG) and the low mortality isolate (VE) exhibited intermediate lesion profiles with very little liver pathology typically observed for ISA. The observed variation in pathology in a standardized infection trial underscores that the progression of ISA disease is influenced by genetic variation among ISAV-HPRΔ isolates, influencing replication capacity, infection dynamics, and tissue tropism.

### Viral load kinetics in blood and tissues

Quantitative PCR analysis of viral RNA in gill, heart, and blood mapped the kinetics of viral replication and dissemination. For all isolates, replication in gills was evident at 1 DPC, confirming local replication in surface epithelium during the primary phase of infection. The viral RNA levels in gills did not fully reflect the viral RNA detection in water, suggesting that other epithelial sites, such as skin, may contribute to the shedding and to the observed differences in replication kinetics. The consistent detection of viral RNA in the heart and in blood at 3 DPC across all isolates demonstrates that HPR-deletions observed in the isolates in this study, allowed systemic dissemination from the primary site of ISAV entry. This was independent of pathotypic segment 5 changes such as Q266L point mutation or an insertion, or as for Å, an atypic point mutation in the HPR-region. The dynamics of viral RNA levels at early (1-3 DPC) time points were similar across isolates, although GL showed a generally lower ability to replicate in gill, in line with previous observations(48). ISAV replication in the gill and heart at 7-14 DPC appeared to be the anatomical site and time point most closely associated with mortality, possibly reflecting the degree of endothelial infection. Seven of the ten isolates with the high viral loads in heart tissue represented high and moderate mortality groups, namely GL, VA, S, H16, Å, B, H20. Seven of the ten isolates with the high viral loads in heart tissue represented high and moderate mortality groups, namely GL, VA, S, H16, Å, B, H20. Despite high viral loads, five of the seven isolates within high and moderate mortality categories showed extensive pathology. The other two isolates, S and H20, displayed typical ISA pathological changes but not to the same extent, with H20 showing the changes much later. In contrast, VE isolate displayed lower viral load in heart (log₁₀ 8.0) and limited pathology, consistent with low mortality indicating lower virulence. The persistence of viral RNA in several tissues up to 42 DPC or until last fish was sampled in that group suggests that the infection is not eliminated but maintained even if overt disease is lacking. This is in keeping with clinical field experience where breaking the chain of infection is essential to control ISA. The biological and epidemiological consequences of such persistence could differ between isolates. The temporal correlation of viral loads in blood and tissue further demonstrates that viraemia can be used as a reliable indicator of systemic infection intensity in individual fish. The detection of RBC-bound ISAV HE by flow cytometry complements this finding, supporting that blood plays a role in systemic viral dissemination within the host. Of note, ISAV-RBC binding appeared to be inhibited from 28 DPC, perhaps reflecting infection-induced loss of the ISAV receptor (39).

Collectively, the findings of this study demonstrate pronounced differences in infection kinetics, virus shedding, and disease outcome amongst the recent Norwegian field isolates. This is suggestive of differential virulence. Further elaboration on identifying specific virulence markers would improve vaccination strategies and could allow inclusion of differences in virulence in a risk analysis to tailor outbreak responses and use less extensive measures for example, for low virulent isolates in low-risk situations. However, several factors such as stress due to handling, environmental adverse conditions, genetics and robustness of fish, and efficacy of vaccines influences infection dynamics and disease outcome. Thus, easily applicable virulence markers could complement the routine fish health and welfare surveillance in low mortality ISA outbreaks where culling is postponed. Virulence of a specific isolate may change over time, and thus such approach should be carefully considered before wide application.

### Conclusions and future perspectives

This study demonstrates that ISAV-HPRΔ isolates exhibit a spectrum of virulence, highlighting that HPR-deletions do not alone define interaction between salmon and ISAV-HPRΔ. The use of standardized bath challenge enabled direct comparison among isolates. This revealed intrinsic differences in infection dynamics and virus shedding that likely contribute to the variable severity observed in field outbreaks. Observations from controlled trials may furthermore not reflect the field situation with the variation in genetic background, immune status, and environmental stressors. Future research should focus on identifying molecular determinants underlying this virulence variation, including sequence differences across the genome and their impact on viral entry, replication, shedding and immune evasion. Moreover, integrating virulence markers with pathogen transmission models in the field and host biomarkers could refine risk analysis for ISA outbreaks and guide more proportionate, evidence-based management responses in aquaculture biosecurity management.

## Author Contributions

**Simon Chioma Weli:** Conceptualization, data curation, formal analysis, visualization, writing – original draft, methodology, investigation, project administration, writing – review and editing, funding acquisition, validation. **Sonal Patel:** Conceptualization, data curation, formal analysis, visualization, writing – original draft, methodology, investigation, writing – review and editing, funding acquisition, validation. **Ole Bendik Dale:** Data curation, formal analysis, visualization, writing – original draft, methodology, investigation, writing – review and editing, validation. **Bjørn Spilsberg, Johanna Hol Fosse, and Torfinn Moldal:** Methodology, Data curation, formal analysis, validation, writing – review and editing. **Laura V. Solarte-Murillo and Saima Nasrin Mohammad:** Investigation, writing – review and editing. **Magnus Leithaug, Marit Måsøy Amundsen, Adriana Magalhaes Santos Andresen, and Frieda Betty Ploss:** Investigation, data curation.

## Ethics approval

All animal procedures were approved (FOTS approval FOTS ID 30432) by the Norwegian Food Safety Authority based on the guidelines of the Norwegian Animal Welfare Act.

## Acknowledgements

The authors thank Brit Tørud Henriette Kvalvik, Merete Gåsvær Sture and Britt Saure for providing technical assistance.

## Data Availability Statement

The data that supports the findings from this study are available within reasonable request.

## Accession numbers for sequence data

### Accession segment 5 (F)

**Accession segment 5 (F):**
| No. | Isolates | Acronym | Accession number |
| --- | --- | --- | --- |
| 1. | Glesvaer/2/90 | GL | HQ259675.1 |
| 2. | NO/Heim/70-122-2/2022 | VE | PZ269547.1 |
| 3. | NO/Lofoten/NVI-70-2774/2014 | VA | OQ310960.1 |
| 4. | NO/Heim/NVI-70-150/2022 | HG | PZ269549.1 |
| 5. | NO/Gildeskål/NVI-70-1297/2016 | H16 | PZ269551.1 |
| 6. | NO/Gildeskål/NVI-70-595/2020 | H20 | PZ269553.1 |
| 7. | NO/Salangen/NVI-70-663/2021 | S | PZ269555.1 |
| 8. | NO/Meløy/NVI-70-1169/2015 | Å | PZ269557.1 |
| 9. | NO/Vesteraalen/NVI-70-9227/2013 | B | OQ310956.1 |
| 10. | NO/Nordkapp/NVI-70-72/2020 | L | PZ269545.1 |
| 11. | SK779/06 | SK | EU118819.1 |

### Accession segment 6 (HE)

**Accession segment 6 (HE):**
| No. | Isolates | Acronym | Accession number |
| --- | --- | --- | --- |
| 1. | Glesvaer/2/90 | GL | HQ259676.1 |
| 2. | NO/Heim/70-122-2/2022 | VE | PZ269548.1 |
| 3. | NO/Lofoten/NVI-70-2774/2014 | VA | OQ310972.1 |
| 4. | NO/Heim/NVI-70-150/2022 | HG | PZ269550.1 |
| 5. | NO/Gildeskål/NVI-70-1297/2016 | H16 | PZ269552.1 |
| 6. | NO/Gildeskål/NVI-70-595/2020 | H20 | PZ269554.1 |
| 7. | NO/Salangen/NVI-70-663/2021 | S | PZ269556.1 |
| 8. | NO/Meløy/NVI-70-1169/2015 | Å | PZ269558.1 |
| 9. | NO/Vesteraalen/NVI-70-9227/2013 | B | OQ310968.1 |
| 10. | NO/Nordkapp/NVI-70-72/2020 | L | PZ269546.1 |
| 11. | SK779/06 | SK | EU118820.1 |

